# IL-10/ACOD1 axis regulates catabolism of phagocytosed lipids in trained macrophages

**DOI:** 10.64898/2026.08.18.745517

**Authors:** Mack B. Reynolds, Annalise Bond, Emily M.J. Fennell, Kym J. Grae, Emeline Joulia, Matthew P. Donnelly, Melissa A. Johnson, Aurélie Laguerre, Gladys R. Rojas, Matthew J. Kolar, Janelle S. Ayres, Christian M. Metallo, Gerald S. Shadel

## Abstract

Macrophages clear excess host and microbial debris to restore homeostasis in inflamed tissues, yet the regulation and molecular fate of phagocytosed lipids during innate immune training remains largely unexplored. Leveraging stable isotope tracing of ^13^C-labeled bacteria, we establish an experimental framework to track microbe-to-host lipid transfer and define the fates of microbial lipids in macrophages *in vitro* and *in vivo*. While naïve macrophages scavenge phagocytosed bacterial fatty acids into the host lipidome, TLR4-trained macrophages direct flux to mitochondria for β-oxidation or lipid droplets in the context of mitochondrial dysfunction. While TLR4 signaling increases ACOD1 expression to produce itaconate that throttles TCA flux, trained macrophages produce IL-10 that reduces ACOD1 to sustain bacterial lipid disposal and promote resolution. These findings reveal an IL-10/ACOD1 regulatory axis in trained macrophages that reprograms lipid metabolism to optimally reestablish tissue homeostasis post-inflammation.

## Introduction

Resolution of tissue inflammation depends on the large-scale disposal of self and foreign biomass. Macrophages, a diverse and ubiquitous class of professional phagocytes, occupy this resolving niche in most tissues^1^. Post-inflammation, the clearance of lipid biomass poses unique challenges. First, aberrant lipids arising from the microbial lipidome or modifications of the host lipidome can be highly immunogenic^2–5^. Further, breakdown of these lipids releases free fatty acids that can be toxic if not rapidly esterified^6,7^. Finally, excess lipid deposition can exacerbate tissue damage and inflammation^8,9^. Failure to efficiently clear inflammatory lipids is a hallmark of metabolic diseases such as atherosclerosis and steatosis and autoimmune disease in the form of antiphospholipid syndrome^10,8,9,11–13^. Thus, macrophages must maintain robust lipid catabolic pathways and other tailored solutions to manage phagocytosed lipid burden to maintain tissue homeostasis.

Initial exposure to microbial pathogen-associated molecular patterns (PAMPs) or host damage-associated molecular patterns (DAMPs) recalibrates the innate immune response to subsequent challenges in an antigen-independent process known as innate immune memory^14,15^. Myeloid-specific innate immune memory orchestrates cross-microbe protection following vaccination, tolerance during sepsis, and chronic inflammation in autoimmune disease^15–20^. To date, most studies of innate immune memory have focused on myeloid signaling functions, primarily through epigenetic regulation of cytokine gene expression^21–25^. Nevertheless, it has been appreciated for nearly a decade that pro-inflammatory cytokine responses alone cannot explain the protection against infection conferred by innate immune memory, though clues have emerged that indicate phagocytic and antimicrobial tuning may be key factors^26,27^. Importantly, innate immune training of the essential catabolism of cargo engulfed by phagocytes remains to be defined.

While early radiolabeling studies showed that bacterial biomass could be incorporated into phagocyte lipids^28^, precisely how phagocytosed bacterial lipids are metabolized within macrophages or how inflammatory experience influences their utilization remains unclear. Using liquid chromatography (LC) and gas chromatography (GC) mass spectrometry (MS)-powered stable isotope tracing of ^13^C-labeled bacterial particles, we investigate the flux of phagocytosed bacterial lipids through recycling and degradative pathways within macrophages *in vitro* and *in vivo*. Our results demonstrate that prior Toll-like receptor 4 (TLR4) exposure trains macrophage catabolism of bacterial lipids. We identify the pro-resolution cytokine interleukin 10 (IL-10) to be key in coordinating TLR4-trained catabolism of bacteria-derived fatty acids by dampening production of itaconate, which acts as a throttle of TCA cycle flux. These results highlight fine-tuning of lipid and mitochondrial metabolism as key components of macrophage training that is regulated by IL-10 and itaconate to reestablish tissue homeostasis post-inflammation.

## Results

### TLR4 training drives macrophages toward mitochondrial catabolism versus recycling of bacterial fatty acids

Since innate immune memory of TLR4 engagement is protective against sepsis^18,29^, we reasoned that TLR4 could train pro-resolution phagocytic catabolism. To study trained innate immunity in a well-controlled system, we adopted an established *in vitro* lipopolysaccharide (LPS; TLR4 agonist) activation and recovery training model using primary murine bone marrow-derived macrophages (BMDMs) (**Fig S1A**)^18^. Naïve and trained BMDMs displayed comparable, high phagocytic capacity when fed heat-killed *Escherichia coli* (HKEC) particles (**Fig S1B-C**), allowing assessment of the fate of phagocytosed bacterial lipids to be determined without confounding effects of differential uptake. To assess catabolism of phagocytosed lipid cargo at the molecular level, we generated stable isotope-labeled HKEC by growth in minimal media with ^13^C_6_ glucose as the sole lipogenic carbon source. As expected, the major lipid class present in ^13^C-labeled (^13^C) HKEC was phosphatidylethanolamine (PE), with a fatty acid profile dominated by 16-18 carbon length straight chain and monounsaturated fatty acids as well as corresponding cyclopropane fatty acids that are uniquely synthesized by bacteria (**Fig S2A**)^30,31^. The entire ^13^C HKEC PE pool was labeled with most of the pool being uniformly ^13^C labeled (**Fig S2B**). The ^13^C HKEC were then fed to macrophages and scavenging, recycling, and catabolism of bacterial lipids were monitored using stable isotope tracing coupled with LC-MS lipidomics (**Fig 1A-D**). Importantly, bacterial lipids were distinguished from host lipids based on expected isotopic labeling, and U^13^C PE could be further validated by detection of a M+2 label in the characteristic phosphoethanolamine headgroup neutral loss ion in MS^2^ (**Fig S2C-D**). Despite normal initial bacterial uptake, TLR4-trained BMDMs displayed improved degradation of bacterial lipids as early as 6 h post-stimulation (**Fig 1A**) and heightened clearance of bacterial fatty acids by 24 h post-stimulation (**Fig 1B**). This increased clearance of bacterial fatty acids could either result from recycling into the host lipidome or transport to mitochondria or peroxisomes for oxidation. To investigate fatty acid utilization in greater detail, we leveraged the observation that the majority of *E. coli* fatty acid is palmitate (C16:0) with a large portion as palmitoleate (C16:1), resulting in a distinct isotopic enrichment (M+16) within the host lipidome upon recycling (**Fig 1C Fig S2E-F**).

**Fig 1.**
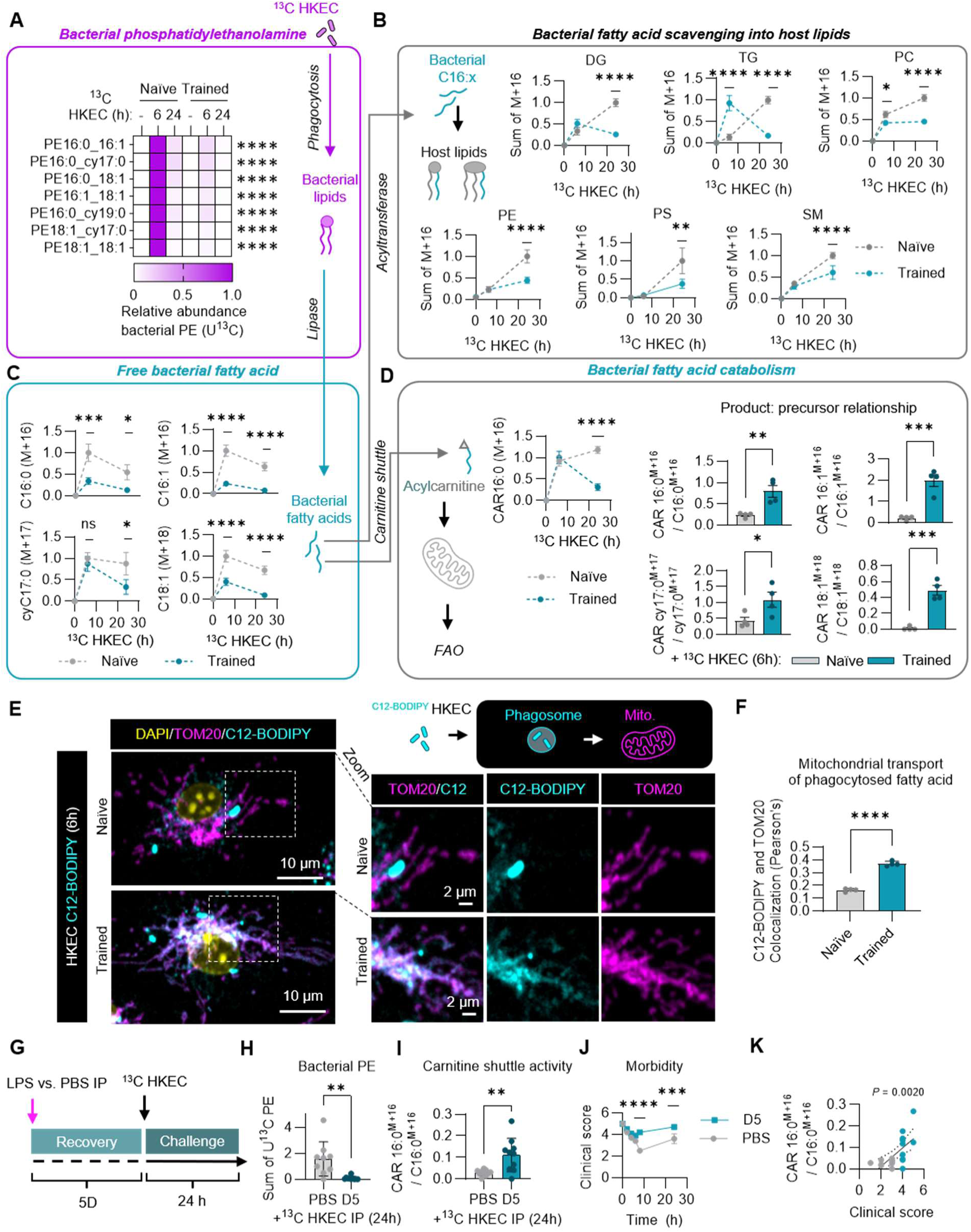
TLR4 trains mitochondrial catabolism of phagocytosed bacterial lipids by macrophages. Naïve or TLR4-trained BMDMs were stimulated ± ^13^C heat-killed *E. coli* (HKEC) for 6 or 24 h and levels of bacteria-derived U^13^C phosphatidylethanolamine (**A**) and free fatty acids (**B**) were monitored by stable isotope tracing high-resolution liquid chromatography mass spectrometry (LC-MS). **C.** LC-MS analysis of bacterial fatty acid recycling into the host lipidome compared between naïve or trained BMDMs stimulated ± ^13^C HKEC for 6 or 24 h and measured as the relative abundance of summed M+16 isotopologues of MS^2^ verified lipids within each lipid class, diacylglycerol (DG), triacylglycerol (TG), phosphatidylcholine (PC), phosphatidylethanolamine (PE), phosphatidylserine (PS), and sphingomyelin (SM). **D.** LC-MS analysis of bacterial fatty acid catabolism via the carnitine shuttle, measured as either kinetic analysis of the relative abundance of ^13^C bacterial fatty acid-loaded acylcarnitine (M+16 for palmitoylcarnitine shown) or the product:precursor ratio of bacterial fatty acid-loaded acylcarnitine (CAR) relative to the ^13^C bacteria-derived free fatty acid abundance at 6 h post-stimulation. **E.** Representative maximum intensity projection of Z-stack confocal micrographs of naïve and trained BMDMs fed with C12-BODIPY-loaded HKEC for 6h and analyzed by immunofluorescence assay against TOM20 with DAPI counterstain. **F.** Average Pearson’s correlation coefficient between C12-BODIPY and TOM20, calculated per cell for naïve and trained BMDMs. **G.** Experimental design of *in vivo* assessment of bacterial lipid catabolism D5 post-sublethal endotoxemia recovery or PBS control rechallenged with IP injection of ^13^C HKEC particles. LC-MS analysis of summed bacteria-derived (U^13^C) PE (**H**) and the ratio of bacterial fatty acid-loaded palmitoylcarnitine (M+16) to bacteria-derived free palmitate (M+16) from peritoneal cavity exudate (**I**). **J.** Clinical scoring of mouse morbidity compared between PBS and LPS-recovered mice rechallenged with HKEC IP. **K.** Correlation between clinical score at 8 h post-HKEC challenge with carnitine shuttle activity, calculated as the ratio of bacterial fatty acid-loaded palmitoylcarnitine (M+16) to bacteria-derived free palmitate (M+16). Graphs represent the mean of n = 4 biological replicates with SEM error bars. *p*-values were calculated with an unpaired *t*-test for single comparisons (**D**, **F, H**, **I**), two-way ANOVA with Tukey’s post-test for grouped analysis (**A-D** and **J-K**), and linear regression for correlation analysis (**K**). The *p*-values displayed are adjusted for multiple comparisons. ns > 0.05; ∗*p* < 0.05; ∗∗*p* < 0.01; ∗∗∗*p* < 0.001; ∗∗∗∗*p* < 0.0001

While multiple forms of bacterial lipid salvage could be detected, such as bacterial lysoPE reacylation and elongation of host fatty acid with bacteria-derived acetyl-CoA, the major isotopic pattern across lipid classes was this M+16 whole fatty acid recycling pattern (**Fig S2F**). We analyzed this isotopic pattern kinetically in naïve or TLR4-trained BMDMs across a panel of major host lipids, diacylglycerols (DG), triacylglycerols (TG), phosphatidylcholines (PC), PE, phosphatidylserines (PS), and sphingomyelins (SM). Interestingly, naïve macrophages tended to recycle bacterial fatty acids into the host lipidome, whereas trained macrophages had reduced incorporation of bacterial fatty acids into all lipid classes by 24 h post stimulation. In line with this observation, we noted that relative conversion of bacterial fatty acids, palmitate (C16:0), palmitoleate (C16:1), vaccenate (C18:1), and cyclopropane C17:0 (cyC17:0) into acylcarnitines for mitochondrial transport and consumption was higher in trained macrophages (**Fig 1D**, **Fig S3A**). Thus, we reasoned that mitochondrial β-oxidation could be the major pathway for disposal that reduces bacterial fatty acid incorporation into the host lipidome.

In a complementary approach, we spatially tracked bacterial fatty acids in macrophages to assess mitochondrial transport efficiency. To achieve this, we loaded HKEC with the fluorescent fatty acid analog C12-BODIPY, which is metabolized and trafficked analogously to endogenous long-chain fatty acids^32^. We observed that C12-BODIPY-loaded HKEC was well retained within bacterial particles at the macroscopic and microscopic levels (**Fig S3B-C**). Consistent with our stable isotope tracing data, we observed increased transport of bacteria-derived C12-BODIPY away from bacterial particles (**Fig S3C-D**) and into mitochondria in trained macrophages compared to naïve macrophages (**Fig 1E-F**). As an added control to validate that C12-BODIPY accurately reflects mitochondrial fatty acid trafficking, we employed differential permeabilization with either digitonin, which permeabilizes the plasma membrane and mitochondrial outer membrane, or Triton-X100, which permeabilizes all membranes. We found that mitochondrial C12-BODIPY signal was retained after digitonin but not after Triton-X100 permeabilization, consistent with localization to the mitochondrial matrix, the known site of β-oxidation (**Fig S3E-G**).

### Recovery from sepsis trains resolution-phase macrophage lipid catabolism

We next sought to investigate whether the lipid catabolic training we observed *in vitro* occurred in an *in vivo* context. We adopted a sublethal endotoxemia sepsis model, whereby mice were injected intraperitoneally with 2 mg/kg LPS or PBS control and monitored for 1, 3, 5, or 7 days post-injection (**Fig S4A**). Luminex-based cytokine analysis revealed peak systemic cytokine production around the first day of LPS injection, with none of the cytokines detectable above background by the fifth day post-stimulation (**Fig S4B**). The transient burst of inflammatory cytokine production matched closely our *in vitro* trained immunity model, giving discrete naïve (PBS), acute inflammatory (D1), and resolution (D3+) phases. Peritoneal macrophages were harvested at each timepoint and challenged with either ^13^C HKEC or C12-BODIPY-loaded HKEC and analyzed by LC-MS and high-content confocal microscopy, respectively. Consistent with our *in vitro* TLR4 training observations (**Fig 1A-D**), we found that by 5 days post-recovery from sublethal endotoxemia, peritoneal macrophages exhibited enhanced bacterial phospholipid catabolism and increased bacterial fatty acid transport to mitochondria via the carnitine shuttle (**Fig S4C-D**). We visualized a similar phenomenon when tracing C12-BODIPY-loaded HKEC with confocal microscopy, whereby resolution-phase peritoneal macrophages more efficiently shuttled bacteria-derived C12-BODIPY into mitochondria (**Fig S4E-F**). Thus, systemic inflammation in a murine model trains resolution-phase phagocytosed lipid catabolism by macrophages.

Next, we compared *in vivo* catabolism of bacterial biomass between naïve (PBS) and resolution-phase (D5 post-LPS) mice injected IP with a high dose of ^13^C HKEC (1x 10^10^ CFU equivalents)^33,34^ (**Fig 1G**). LC-MS analysis of peritoneal exudates from these mice revealed a striking improvement in bacterial phospholipid catabolism and elevated carnitine shuttling of bacterial fatty acid (**Fig 1H-I**), in line with TLR4 training of macrophages *in vitro* (**Fig 1A-D**). Overall, we found that resolution-phase mice handled bacterial challenge better than naïve mice, as indicated by improved clinical and health scores, reduced weight changes, and stable temperature readings (**Fig 1J Fig S4G**), consistent with prior studies^29^. Finally, we correlated health outcomes with carnitine shuttling of bacteria-derived fatty acid and observed that carnitine shuttle activity correlated best with health scores and clinical scores, suggesting that enhanced lipid catabolism may confer protection during bacterial sepsis (**Fig 1K Fig S4H**).From these *ex vivo* and *in vivo* studies, we conclude that heightened macrophage lipid catabolic capacity is programmed after training post-endotoxemia to support restoration of lipid homeostasis.

### Mitochondrial dysfunction drives storage of exogenous fatty acids in lipid droplets within TLR4-trained macrophages

Post-inflammation, resolution-phase macrophages adopt a fatty acid oxidation (FAO)-driven mitochondrial metabolic state, which depends on carnitine shuttle-fed long-chain fatty acid β-oxidation in mitochondria^35^. Since we observed that bacterial fatty acids were preferentially shuttled into mitochondria, we investigated whether this could be mediated by a global increase in carnitine shuttle activity. Therefore, we measured the abundance of long and short chain acylcarnitines by LC-MS. We observed that trained BMDMs had elevated levels of both long-chain and short-chain acylcarnitines, indicative of accelerated carnitine shuttle-coupled FAO, whereas continuously activated macrophages only had elevation of short chain acylcarnitines (**Fig S5A**). We predicted that this increase in FAO would enhance oxidative phosphorylation (OXPHOS) in trained BMDMs. To test this, we performed Seahorse extracellular flux analysis of naïve and trained BMDMs treated with the carnitine palmitoyltransferase 1A (CPT1A) inhibitor etomoxir or vehicle control (**Fig S5B**). We found that trained BMDMs exhibited elevated basal mitochondrial respiration (oxygen consumption rate; OCR), which was reduced by etomoxir treatment (**Fig S5C**). Therefore, we conclude that increased OXPHOS in TLR4-trained macrophages is supported by elevated FAO.

Based on the observation that phagocytosed bacterial fatty acids were better loaded onto acylcarnitines and consumed faster in trained macrophages (**Fig 1D**), we reasoned that phagocytosed lipid catabolism would depend on the carnitine shuttle and enhanced FAO-coupled OXPHOS. Thus, we treated ^13^C HKEC-fed naïve and trained BMDMs with the CPT1A inhibitor etomoxir, the mitochondrial Complex III inhibitor antimycin A, or vehicle control and measured the abundance of bacteria-derived phospholipids (**Fig 2A**), fatty acids (**Fig 2B**), and bacterial fatty acid-loaded acylcarnitine (**Fig 2C**). OXPHOS inhibition impaired bacterial phospholipid degradation and free fatty acid clearance in naïve BMDMs, yet trained BMDMs retained a mechanism to sequester bacteria-derived lipids post-catabolism. Using stable isotope tracing and high-resolution LC-MS-based lipidomics, we measured lipids with bacterial C16:x fatty acid incorporation by tracking M+16 isotopic labeling signatures. The major destination of bacterial fatty acids diverted from mitochondrial FAO was neutral lipid stores, specifically triacylglycerols (TGs) (**Fig 2D-E**). Measuring the relative abundances of the corresponding host-derived lipids (M+0), we observed a similar increase in TG pools post FAO-coupled OXPHOS blockade in trained BMDMs, highlighting the interplay between mitochondrial function and lipid accumulation (**Fig S5D**). Thus, trained macrophages are capable of efficiently storing fatty acids derived from phagocytosed cargo when their catabolic disposal route is compromised.

**Fig 2.**
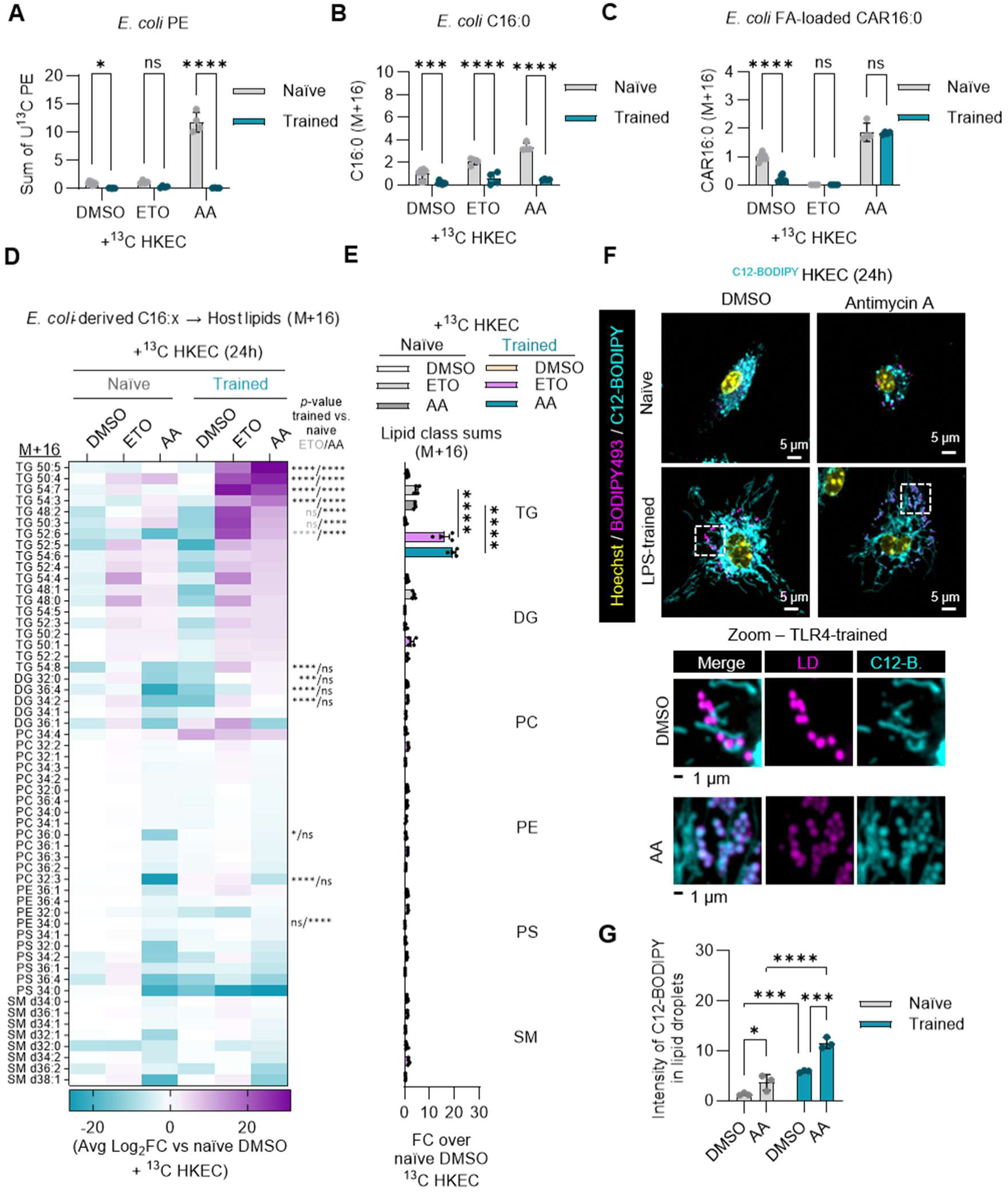
Elevated fatty acid storage capacity accommodates trained bacterial lipid catabolism under conditions of OXPHOS blockade. LC-MS analysis of summed bacteria-derived (U^13^C) PE (**A**) bacteria-derived (U^13^C) palmitate (**B**), and bacterial fatty acid-loaded palmitoylcarnitine (M+16) (**C**) from naïve or trained BMDMs rechallenged with ^13^C HKEC for 24 h in the presence of etomoxir (ETO), antimycin A (AA), or vehicle control (DMSO). **D.** LC-MS analysis of bacterial fatty acid recycling into the host lipidome compared between naïve or trained BMDMs stimulated with ^13^C HKEC for 24 h in the presence of ETO, AA, or DMSO and measured as the Log_2_ fold change over naïve BMDMs treated with vehicle control and ^13^C HKEC of relative abundance of individual M+16 isotopologues of MS^2^ verified lipids within each lipid class, diacylglycerol (DG), triacylglycerol (TG), phosphatidylcholine (PC), phosphatidylethanolamine (PE), phosphatidylserine (PS), and sphingomyelin (SM). **E.** LC-MS stable isotope analysis as presented in **D**, except relative abundance of summed M+16 isotopologues is presented per condition. **F.** Representative maximum intensity projection of Z-stack confocal micrographs of naïve and trained BMDMs fed with C12-BODIPY-loaded HKEC for 24 h in the presence of AA or DMSO and analyzed by live cell imaging with lipid droplet counterstain BODIPY493 and nuclear counterstain Hoechst. **G.** Quantification of the average integrated intensity of C12-BODIPY within BODIPY493-segmented lipid droplets per cell. Graphs represent the mean of n ≥ 3 biological replicates with SEM error bars for all graphs. *p*-values were calculated using two-way ANOVA with Sidak’s post-test ns > 0.05; ∗*p* < 0.05; ∗∗*p* < 0.01; ∗∗∗*p* < 0.001; ∗∗∗∗*p* < 0.0001.

In *E. coli*, the primary C16 fatty acids, palmitic acid (C16:0) and palmitoleic acid (C16:1), are also common to mammalian cells. Thus, we next sought to identify if similar mechanisms were in place for the trained catabolism of fatty acids that are more abundant in bacterial systems such as cyclopropane and branched-chain fatty acids (**Fig S6A-F**, **Fig S7A**). To assess this, we measured lipidomic enrichment of ^13^C HKEC-derived M+18 (vaccenic acid) and M+17 (cyclopropane C17:0 fatty acid), and U^13^C glucose-fed heat-killed *Staphylococcus aureus* (HKSA)-derived M+10 (*anteiso*-C15:0 monomethyl branched-chain fatty acid, derived from elongation of ^12^C isoleucine)^36^. Inhibition of mitochondrial Complex III led to the accumulation of these bacterial free fatty acids in naïve BMDMs but to a lesser degree in trained BMDMs, indicating sustained, trained utilization despite a sharp increase in bacterial fatty acid-loaded acylcarnitines following OXPHOS blockade (**Fig S7B-C**). More broadly surveying the lipidome, we observed that the major destination of bacterial-specific fatty acids in trained macrophages post-Complex III inhibition was TG pools (**Fig S7D**). We conclude that neutral lipid stores are adapted in trained macrophages as a reservoir for excess, diverse phagocytosed lipid-derived fatty acids. Importantly, bacterial fatty acids were enriched within the TG pool but not other major lipid pools, like PC, PE, or SM in trained macrophages, following Complex III inhibition, highlighting a potential function of lipid droplets in foreign lipid sequestration. To spatially track the fate of bacterial lipids, we used C12-BODIPY-loaded HKEC coupled to live imaging confocal microscopy analysis of trained macrophages following OXPHOS disruption via Complex III inhibition (**Fig 2F-G**). Consistent with our flux and lipidomics data, we observed that bacteria-derived C12-BODIPY was diverted to lipid droplets, the major site of neutral lipid storage. These results indicate that trained macrophages have enhanced OXPHOS-coupled FAO. Interruption of this catabolic pathway at the points of mitochondrial entry via the carnitine shuttle or OXPHOS-coupled FAO results in substantial storage of phagocytosed fatty acids within neutral lipid pools.

### ACOD1 and itaconate throttle TCA cycle-mediated lipid disposal in trained macrophage

Fatty acid oxidation is tightly linked to cellular bioenergetics, producing acetyl-CoA that is fed into the TCA cycle to yield reducing equivalents and release carbon as CO_2_. However, release of carbon through this oxidative mechanism depends on an intact TCA cycle with capacity for continuous cycling. Notably, TLR4 signaling drives macrophages to produce the immunometabolite itaconate, which blunts TCA cycle flux by inhibiting succinate oxidation to fumarate by succinate dehydrogenase (SDH) / mitochondrial Complex II^37,38^. Critically, we observed that the transcript of *Acod1,* which encodes the itaconate-producing enzyme ACOD1, was the most highly induced transcript encoding a mitochondrial protein in trained compared to naïve macrophages (**Fig 3A**), which we confirmed resulted in more ACOD1 protein by immunoblot (**Fig 3B**). Indeed, we observed that, even as the production of the pro-inflammatory cytokine tumor necrosis factor (TNF) subsided in trained BMDMs (**Fig S8A**), ACOD1 protein expression remained high (**Fig S8B**). Furthermore, we measured steady-state itaconate and succinate levels in a targeted metabolomics GC-MS platform and observed a robust increase in itaconate and a corresponding increase in succinate levels in trained BMDMs (**Fig 3C**). Finally, we observed that TLR4-trained macrophages produced elevated levels of intracellular and extracellular itaconate (**Fig S8C**). Because normal TCA cycling is reduced by SDH/Complex II inhibition, our data suggest that itaconate secretion may alleviate TCA cycle load in trained macrophages.

**Fig 3.**
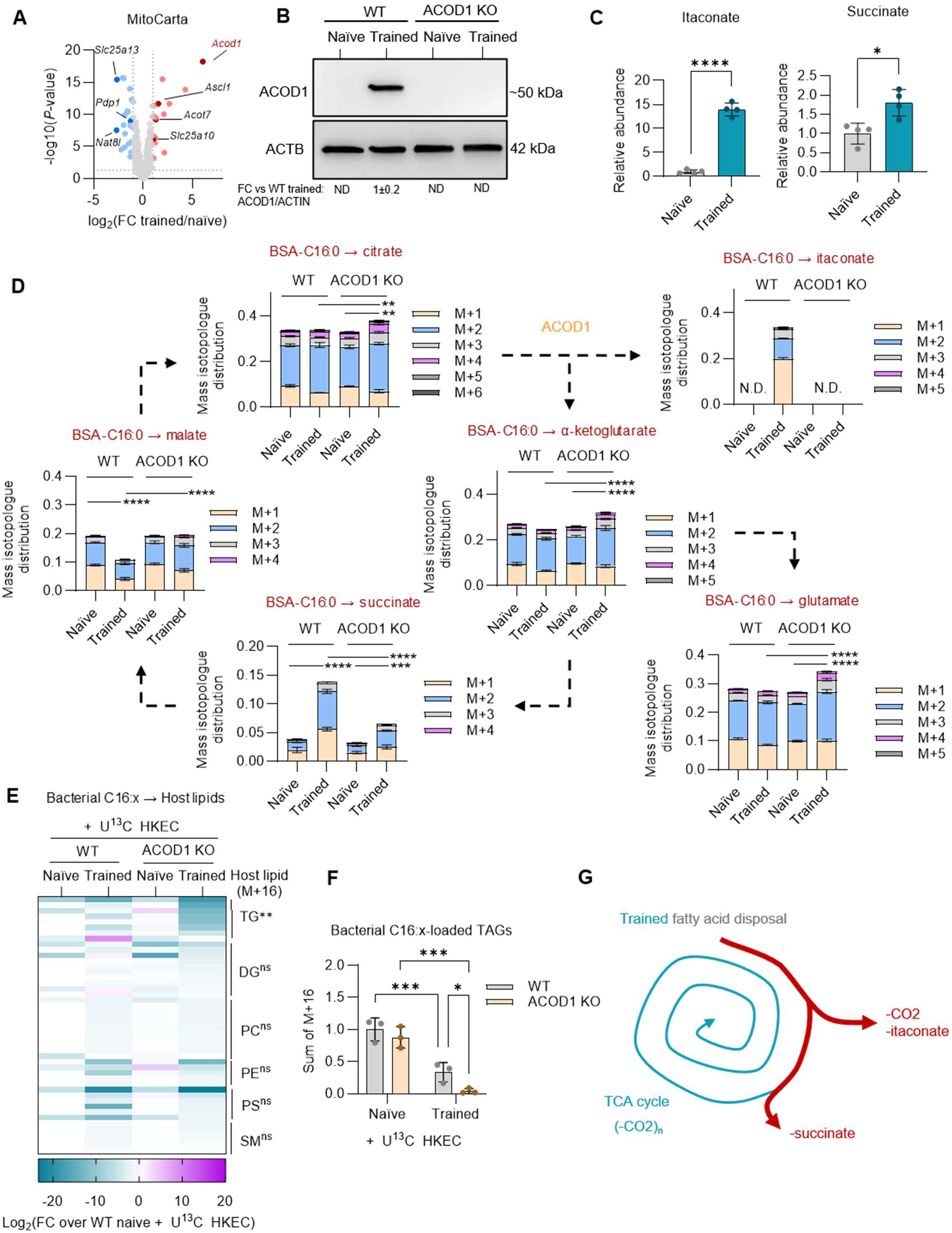
Itaconate throttles fatty acid-derived TCA cycle flux in TLR4-trained macrophages. **A.** Volcano plot of RNA sequencing analysis of trained vs. naïve BMDMs, filtered for MitoCarta3.0 annotated transcripts, which encode mitochondria-localized proteins. TCA cycle and lipid metabolism-related genes are labeled. **B.** Western blot analysis of ACOD1 expression in WT and ACOD1 KO naïve and trained BMDMs with β-actin (ACTB) as a loading control. **C.** Gas chromatography mass spectrometry (GC-MS) analysis of itaconate and succinate in naïve and trained macrophages. **D.** GC-MS-based stable isotope tracing of U^13^C palmitate bound to bovine serum albumin (BSA) for 4 h into TCA cycle intermediates, reporting mass isotopologue distributions of fractions of metabolite pools labeled with the indicated isotopologues compared between WT and ACOD1 KO naïve and trained BMDMs. **E.** LC-MS analysis of bacterial fatty acid recycling into the host lipidome compared between WT and ACOD1 KO naïve and trained BMDMs stimulated with ^13^C HKEC for 24 h and measured as the Log_2_ fold change over naïve HKEC-fed WT BMDMs of relative abundance of individual M+16 isotopologues of MS^2^ verified lipids within each lipid class: diacylglycerol (DG), triacylglycerol (TG), phosphatidylcholine (PC), phosphatidylethanolamine (PE), phosphatidylserine (PS), and sphingomyelin (SM). **F.** LC-MS stable isotope analysis as presented in **E**, except relative abundance of summed M+16 isotopologues across TG species is presented per condition. **G.** Illustration of competing carbon outlets via TCA cycling and the itaconate shunt in TLR4-trained BMDMs. Graphs represent the mean of n ≥ 3 biological replicates with SEM error bars for all graphs. *p*-values were calculated using an unpaired *t*-test (**C**), and two-way ANOVA with Sidak’s post-test (**D-F**). For **D**, *p-*values were calculated based on 1-M0 fractional labeling method. For **E**, statistics refer to WT trained vs. ACOD1 KO trained normalized peak area sums across validated species within each class. ns > 0.05; ∗*p* < 0.05; ∗∗*p* < 0.01; ∗∗∗*p* < 0.001; ∗∗∗∗*p* < 0.0001.

Next, we sought to compare the metabolic fate of oxidized fat acids in naïve versus trained macrophages in the presence or absence of endogenous itaconate. To this end, we generated naïve or trained WT or ACOD1 KO BMDMs (**Fig 3B**) and fed them BSA loaded with U^13^C palmitate and measured TCA cycle intermediate fractional labeling with GC-MS (**Fig 3D**). Within 4 h, increased itaconate and succinate labeling was observed in trained compared to naïve macrophages with no difference or a reduction in other TCA intermediates. This result suggests accumulation of fatty acid-derived carbon into itaconate and succinate pools in trained macrophages. Interestingly, ACOD1 KO increased labeling of other TCA intermediates, such as citrate and alpha-ketoglutarate, including higher-labeled isotopologues, in trained macrophages. Labeling changes generally reflected shifts in TCA cycle intermediate pool sizes, except in the case of citrate which was reduced in total pool size in trained ACOD1 KO BMDMs despite increased fat-derived labeling (**Fig S8D**). Overall, we conclude that itaconate serves to throttle TCA cycle flux in trained macrophages, with itaconate serving as a release mechanism for carbon load. However, it remains unclear whether the itaconate shunt would facilitate or impair the clearance of phagocytosed lipids in trained macrophages. To address this question, we fed ^13^C HKEC to naïve or trained WT or ACOD1 KO BMDMs and monitored bacteria-derived fatty acid recycling into the host lipidome. While WT and ACOD1 KO trained BMDMs both exhibited a general reduction in bacterial fatty acid recycling into the lipidome compared to naïve controls, trained ACOD1 KO had an even greater reduction in storage of bacteria-derived fatty acids into TGs without affecting the pool size of host-derived TGs (**Fig 3E-F**, **Fig S8E**). We conclude that itaconate throttles phagocytosed bacterial fatty acid oxidation by inhibiting TCA cycle flux through SDH/Complex II, suggesting its production may serve to attenuate or control excessive catabolic TCA cycle flux in trained macrophages (**Fig 3G**).

### IL-10 suppresses ACOD1 expression to regulate bacterial fatty acid catabolism

Inflammation is a highly dynamic process which requires the continuous remodeling of the tissue microenvironment by recruited immune cells. Since in a confined infection or inflammatory event, immune cells would not sense microbial cues directly prior to recruitment, we hypothesized that trained lipid catabolic programming could be conferred to peripheral myeloid cells via systemic cytokines. While the induction of certain pro-inflammatory cytokines is suppressed post-recovery from TLR4 activation^18^, other cytokines may be enhanced and help to shape the lipid catabolic response of local and recruited macrophages. To assess this potential mechanism, we performed Luminex-based secreted cytokine profiling, comparing naïve and trained BMDMs stimulated with HKEC for 24 h. While pro-inflammatory cytokines like IFN-β and IL-12 were reduced, other cytokines were enhanced, such as VEGF-A, IL-10, and CXCL1 (**Fig 4A Fig S9A**). We focused on IL-10 because of its known essential function in inflammation resolution, influence on mitochondrial function, and high expression observed in the early inflammatory phase of our sublethal endotoxemia *in vivo* model (**Fig S4B**)^39–41^. Therefore, we hypothesized that IL-10 is a key component of the TLR4-trained response in macrophages that enhances catabolism of phagocytosed bacterial lipid. To test this, we pre-treated BMDMs for 24 h with or without IL-10 and then stimulated these cells for 24 h with HKEC. Using a Seahorse XF assay, we determined that macrophages exposed to IL-10 exhibited increased basal respiration during HKEC stimulation, consistent with prior work (**Fig S9B-C**)^39^. In parallel, we fed naïve or IL-10-treated BMDMs ^13^C HKEC and measured the incorporation of bacterial fatty acids into the host lipidome using LC-MS. We observed that IL-10-treated macrophage had reduced accumulation of bacterial fatty acids in TGs (**Fig S9D**). Thus, IL-10 is sufficient to enhance catabolism and clearance of phagocytosed bacterial fatty acid.

**Fig 4.**
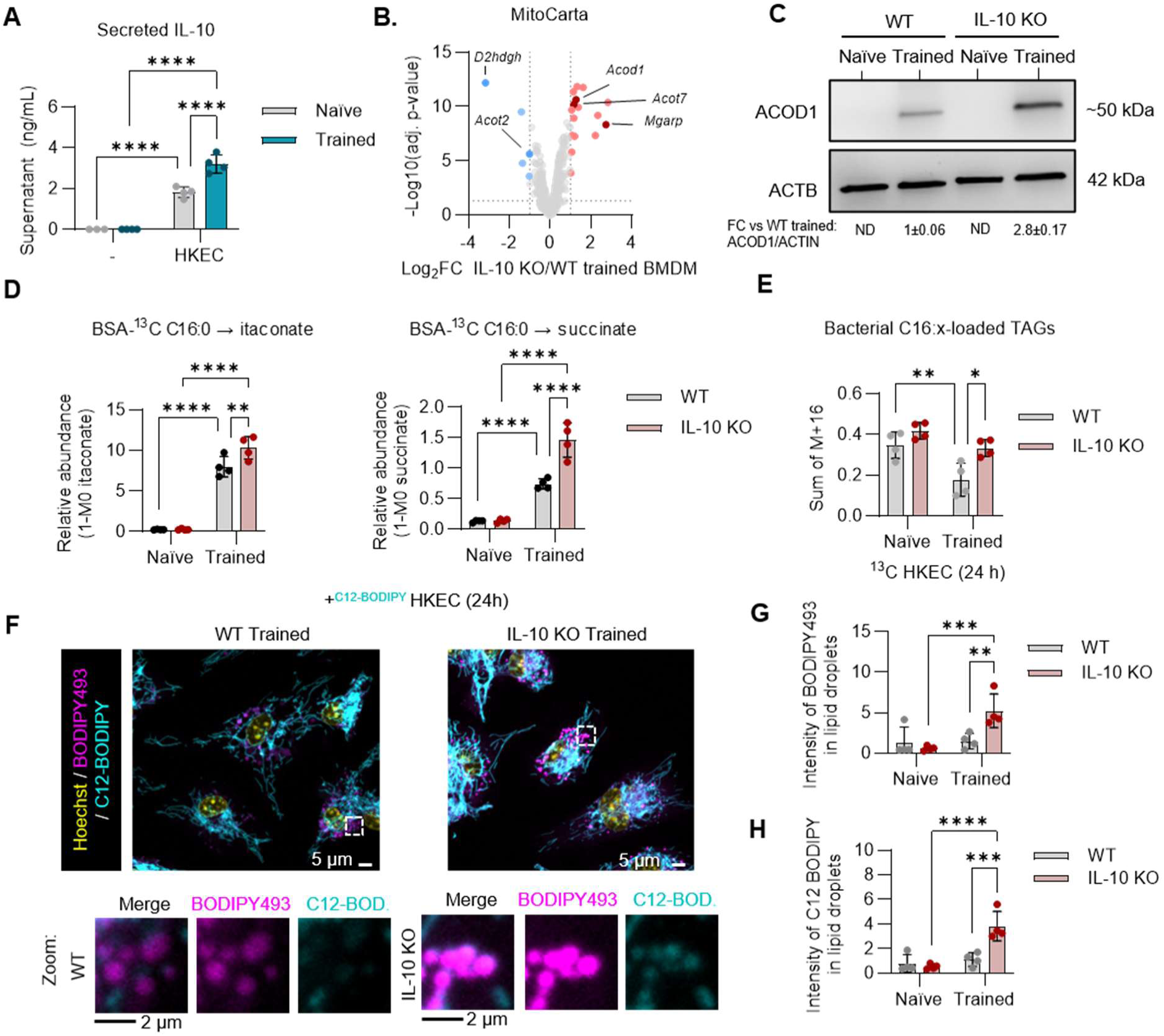
IL-10 constrains *Acod1* to promote phagocytosed lipid catabolism in trained macrophages. **A.** Luminex detection of secreted IL-10 from naïve or trained BMDMs stimulated with HKEC or without (-) for 24 h. **B.** Volcano plot of RNA sequencing analysis of IL-10 KO vs WT trained BMDMs, filtered for MitoCarta3.0 annotated transcripts. TCA cycle and lipid metabolism-related genes are labeled. **C.** Western blot analysis of ACOD1 expression in WT and IL-10 KO naïve and trained BMDMs with β-actin (ACTB) as a loading control. **D.** GC-MS-based stable isotope tracing of U^13^C palmitate bound to bovine serum albumin (BSA) for 4 h into itaconate and succinate, reporting the relative abundance of the total labeled fraction (1-M0) between WT and IL-10 KO naïve and trained BMDMs. **E.** LC-MS analysis of bacterial fatty acid recycling into the host lipidome compared between WT and IL-10 KO naïve and trained BMDMs stimulated with ^13^C HKEC for 24 h and measured as the relative abundance over naïve HKEC-fed WT BMDMs of summed M+16 isotopologues across MS^2^ verified triacylglycerol species (TG)**. F.** Representative maximum intensity projection of Z-stack confocal micrographs of WT and IL-10 KO naïve and trained BMDMs fed with C12-BODIPY-loaded HKEC for 24 h and analyzed by live cell imaging with lipid droplet counterstain BODIPY493 and nuclear counterstain Hoechst. Zoomed regions of interest are included to demonstrate increased localization of bacteria-derived C12-BODIPY within lipid droplets in IL-10 KO trained macrophages fed with C12-BODIPY HKEC. Quantification of the average integrated intensity of BODIPY493 (total neutral lipid) (**G**) and C12-BODIPY (bacteria-derived fatty acid) within BODIPY493-segmented lipid droplets per cell (**H**). Graphs represent the mean of n = 4 biological replicates with SEM error bars for all graphs. *p*-values were calculated using two-way ANOVA with Sidak’s post-test (**A**, **D, E**, **G**, **H**). ns > 0.05; ∗*p* < 0.05; ∗∗*p* < 0.01; ∗∗∗*p* < 0.001; ∗∗∗∗*p* < 0.0001.

Next, we sought to determine if IL-10 was required for TLR4-trained bacterial lipid catabolism. We generated IL-10 KO BMDMs (**Fig S9E**) and performed Seahorse XF analysis and determined that the TLR4-trained increase in respiration was dependent on IL-10 production (**Fig S9F-G**). We hypothesized that IL-10 might function, in part, to dampen negative regulators of mitochondrial metabolism and enhance TCA cycle flux in the resolution phase of inflammation. To this end, we performed RNA sequencing of TLR4-trained WT or IL-10 KO BMDMs, which revealed that *Acod1* was one of the most suppressed mitochondria-related transcripts by IL-10 (**Fig 4B**). IL-10 also suppressed expression of *Nos2*, indicating that IL-10 suppresses both *Nos2* and *Acod1,* which are the major genes responsible for inhibiting mitochondrial electron transport in inflammatory macrophages (**Fig S9H-I)**^37,42–44^. Furthermore, SDS-PAGE and immunoblot analysis of protein extracts from naïve or TLR4-trained WT or IL-10 KO BMDMs confirmed the role for IL-10 in suppressing ACOD1 protein expression post-training (**Fig 4C**).

Finally, using stable isotope tracing coupled to GC-MS analysis of naïve or TLR4-trained WT or IL-10 KO BMDMs fed U^13^C palmitate-BSA, we observed an elevated accumulation of fat-derived itaconate and succinate in IL-10 KO TLR4-trained BMDMs compared to WT TLR4-trained BMDMs (**Fig 4D**), supporting the notion that IL-10 functions to alleviate fat-derived itaconate and succinate accumulation in trained BMDMs. To assess the contribution of IL-10 to bacterial lipid catabolism, we fed ^13^C HKEC to naïve or TLR4-trained WT or IL-10 KO BMDMs and performed LC-MS analysis of bacterial fatty acid incorporation into the host lipidome. We observed that the absence of IL-10 prevented the TLR4-trained reduction in the storage of bacterial fatty acids into TG pools (**Fig 4E**). In a complementary approach, we performed live-cell confocal imaging of C12-BODIPY-loaded HKEC fed to naïve or TLR4-trained WT or IL-10 KO BMDMs (**Fig 4F**). We found that there was a general increase in total lipid droplet staining intensity based on BODIPY493 as well as bacteria-derived C12-BODIPY incorporation in IL-10 KO trained BMDMs compared to WT trained BMDMs (**Fig 4G-H**). Collectively, these studies identify a regulatory node comprising IL-10, ACOD1, and mitochondrial metabolism which controls lipid clearance by trained macrophages.

## Discussion

Innate immune receptors initiate an inflammatory trajectory that begins with pro-inflammation and ends with pro-resolution innate immune function^1^. Notably, microbial pattern recognition is essential for this resolving transition in wound healing and infection^45^. The resolution phase of inflammation requires coordinated tissue remodeling and metabolic adaptation, including substantial catabolism of self and foreign biomass by phagocytes^46^. While tissue-resident and recruited macrophages are essential arms of this catabolic response, we do not yet understand how inflammatory experience trains this catabolism, especially regarding lipid homeostasis. Lipids are highly dynamic molecular species, which undergo continuous and rapid fatty acyl modification, recycling, remodeling, scavenging, and catabolism^47^. In macrophages, the lipidomic fates of exogenous fatty acids depend on the inflammatory status^48^. Accordingly, the metabolism of phagocytosed cargo is distinct, as demonstrated by a recent study, in which Lesbatz *et al.* employ stable isotope tracing to uncover a unique fate for phagocytosed bacteria-derived polar metabolites^49^. Here, we compared how naïve acutely activated macrophages and post-inflammatory trained macrophages handle phagocytosed bacterial lipids. We demonstrate *in vitro* and *in vivo* that naïve macrophages tend to recycle phagocytosed bacterial lipids, while trained macrophages prefer to shuttle these lipids into catabolic pathways. We propose several possible explanations for how this shift in lipid metabolism could be evolutionarily advantageous post-inflammation. First, inflammation is associated with excessive lipid accumulation in tissue, and this trained catabolic function would promote clearance of debris. Second, removal of potentially immunogenic lipids would help to limit autoimmunity and chronic inflammation. And third, naïve macrophages, under homeostatic conditions, may favor recycling to reduce energy expenditures associated with fatty acid synthesis.

In mice and humans, ACOD1 is robustly induced by macrophages during inflammation to produce the TCA cycle-derived metabolite itaconate, which exhibits immunomodulatory and antimicrobial activity^37,38,50–52^. Itaconate directly inhibits SDH/Complex II and methylmalonyl-CoA mutase, both of which control TCA cycle flux^37,53^. Here, we find that macrophages retain ACOD1 expression post-recovery from inflammatory activation, even as the expression of other pro-inflammatory genes is reduced. This higher basal level of itaconate throttles the accelerated fatty acid flux through the TCA cycle present in trained macrophages. That is, instead of repeated TCA cycling and complete oxidation to CO_2_, post-inflammatory macrophages produce high levels of secreted itaconate. The concept that this itaconate secretory pathway may indeed serve as a carbon removal system downstream of high rates of lipid catabolism is supported by a recent study in which urinary excretion was identified as the primary disposal route for peripheral exogenously delivered itaconate^54^. Here, we demonstrate that ACOD1 KO in trained macrophages results in improved TCA cycle flux from fatty acid and reduced phagocytosed fatty acid storage. Therefore, while itaconate is an effective outlet for fat-derived carbon release from the TCA cycle, itaconate secretion is less efficient than continuous TCA cycle flux in disposing of bacteria-derived fatty acids.

We hypothesized that peripheral cytokines produced during inflammation could train innate immune cells towards increased catabolism in the resolution phase of inflammation without the requirement for direct PAMP or DAMP engagement. Notably, IL-10 is ubiquitously expressed as an anti-inflammatory, pro-resolution signal across virtually all forms of inflammation with multiple redundant signaling pathways converging on its expression^55,56^. IL-10 sustains oxidative phosphorylation and coordinates lipidomic remodeling in inflammatory macrophages^39,40^. Of critical importance, IL-10 is induced by efferocytosis and promotes macrophage lipid uptake and survival in a lipid-laden state^41,57^. Our finding that IL-10 coordinates innate immune training of phagocytosed bacterial lipid catabolism expands our appreciation of IL-10 in resolution-phase macrophage function. We identify that this IL-10-preserved trained macrophage lipid catabolic function works in part due to repression of *Acod1* and itaconate production. We speculate that itaconate inhibits TCA cycle flux during the acute inflammatory phase, and IL-10 negatively regulates *Acod1* and itaconate to restore TCA cycle flux during the resolution phase to support lipid catabolism. Still, IL-10 could enhance other dimensions of this catabolic response by modulating gene expression in other mitochondrial and lipid metabolic pathways.

Lipid accumulation has been connected to chronic inflammatory conditions with serious implications for human health including medical conditions such as steatosis and atherosclerosis^8,9^. A hallmark of inflammatory disease is the formation of lipid-laden “foamy” macrophages, which are considered major contributors to disruptive lipid deposition within tissue^8^. Here, we identified enhancement of TLR4-trained macrophage lipid catabolism post-inflammation which supports clearance of phagocytosed lipids, and yet OXPHOS inhibition led to a massive storage of phagocytosed fatty acids within TGs in lipid droplets. Critically, enhanced phagocytosed fatty acid storage in trained macrophages following OXPHOS blockade extended to common bacterial fatty acids, such as *anteiso* branched-chain C15:0 fatty acid from *S. aureus* and cyclopropane fatty acid C17:0 from *E. coli*. We hypothesize that, under normal conditions, this process functions as a backup mechanism to handle fatty acid overload derived from excess lipid uptake, whereby surplus fatty acids are sequestered within lipid droplet TG pools for later oxidation. However, we predict that this enhanced lipid catabolic pathway in macrophages is particularly susceptible to mitochondrial perturbations, which are common in primary mitochondrial diseases, aging, and inflammation^58,59^. Thus, strategies to preserve mitochondrial function in trained macrophages to restore pro-resolution catabolism without risk of lipid deposition could be of therapeutic value. Collectively, our results identify an IL-10 and ACOD1-regulated metabolic program in trained macrophages that is responsible for the uptake, processing, and catabolism of phagocytosed lipids.

## Acknowledgments

We thank members of the Shadel and Metallo laboratories for helpful discussions. This work was supported by funds from the Salk Institute of Biological Sciences, the Allen Institute and the Spruance Foundation II to G.S.S, who is also the Audrey Geisel Chair of Biomedical Science, the Howard Hughes Medical Institute (J.S.A.), the Salk NCI Cancer Center CCSG P30CA014195 (C.M.M.), and California Institute of Regenerative Medicine grant DISC4-19271 (C.M.M.). M.B.R. was supported by the George E. Hewitt Foundation for Medical Research Postdoctoral Fellowship, M.J.K. was supported by the NIH Director’s Early Independence Award (DP5OD039470), M.P.D. was supported by the NIH grant F30HL178290-01, M.A.J. was supported by NIH grant F31CA278581-03, and A.B. is a Fellow of The Jane Coffin Childs Fund for Medical Research. This work was also supported by the Flow Cytometry Core Facility of the Salk Institute (RRID:SCR_014839) with funding from NCI CCSG: P30 AG068635, the Waitt Advanced Biophotonics Core Facility of the Salk Institute (RRID:SCR_014838), and additional microscopy instrumentation resources at the UC San Diego Agilent Center of Excellence in Cellular Intelligence.

## Author contributions

C.M.M, G.S.S., and M.B.R conceived the study. M.B.R., A.B., E.M.J.F., K.J.G, E.J., M.P.D., M.A.J., A.L., G.R.R., and M.J.K. performed experiments. M.B.R. and A.B. analyzed the data. J.A.S. provided resources and contributed to study design. M.B.R. wrote the original draft, and C.M.M. and G.S.S. reviewed and edited the manuscript with input from all authors. G.S.S. and C.M.M. supervised the study, and C.M.M and G.S.S. acquired funding.

## Materials and Methods

### Mice

Wild-type C57BL/6J (strain #000664; WT), C57BL/NJ-Acod1*^em1(IMPC)J^*/J (strain #029340; Acod1 KO), and B6.129P2-*Il10^tm1cgn^*/J (strain #002251; IL10 KO) mice were purchased from The Jackson Laboratory. Male and female mice aged 8–16 wks were used for both *in vitro* and *in vivo* studies. No sex differences were observed; therefore, data are pooled. All animal procedures were approved by and performed in accordance with the Institutional Animal Care and Use Committee (IACUC) at Salk Institute for Biological Studies.

### Murine bone marrow-derived macrophage (BMDM) differentiation

BMDMs were differentiated according to established protocols^60^. Briefly, femurs and tibiae were isolated, bone marrow was flushed with sterile PBS, and cells were strained through a 40-μm filter to generate a single cell suspension. Bone marrow cells were differentiated to macrophages in bone marrow macrophage media (BMM): complete Dulbecco’s modified eagle medium (DMEM; Gibco, cat. no. 11320033) supplemented with 20% heat-inactivated fetal bovine serum (FBS; Avantor® Seradigm cat. no. 76419-584), 30% L929 cell-conditioned medium, penicillin-streptomycin (100 U/mL), and tissue culture grade beta-mercaptoethanol (50 μM). BMDM differentiation occurred over 6 days, supplementing fresh BMM at day 3 post-isolation. Successful BMDM differentiation was validated by flow cytometric analysis of F4/80 positivity (Thermofisher, 48-4801-80). BMDMs were cryopreserved at day 6 in BMM supplemented with 10% sterile dimethylsulfoxide (DMSO).

### *In vitro* BMDM training and challenge

BMDMs were resurrected from cryopreservation for 24 h prior to experimentation and treated +/- 100 ng/mL lipopolysaccharide (LPS, O111:B4; Sigma-Aldrich, cat. no. 437627) for 24 h in complete DMEM supplemented with 10% heat-inactivated FBS (D10). After 24 h stimulation, cells were washed, supplied with fresh D10, and allowed to recover for 24 h. Cells that received LPS activation and recovery were designated “trained,” while cells that only received media replacement were designated “naïve.” After recovery, BMDMs were fed heat-killed *E. coli* (HKEC), generated as described below, at a multiplicity of infection (MOI) of 25 for indicated durations without changing media. In a subset of experiments, cells were treated with either etomoxir (ETO; Cayman Chemical, cat. no. 11969), antimycin A (AA; Sigma-Aldrich, cat. no. A8674), or DMSO vehicle control.

### Generation of ^13^C-labeled heat-killed bacterial particles

*E. coli* strains K12 MG1655 and O21:H+ were used for *in vitro* and *in vivo* studies, respectively^34,61^. Single *E coli* colonies were picked and grown overnight (16 h) at 37°C, slanted with 200 rpm agitation, in M9 minimal media prepared in-house from Milli-Q^®^ water, 0.1 mM CaCl_2_, 2mM MgSO_4_, 1:4000 dilution of vitamin supplement (ATCC MD-VS), 6.8 g/L Na_2_HPO_4_, 3 g/L KH_2_PO_4_, 0.5 g/L NaCl, and 1g/L NH_4_Cl. *S. aureus* strain USA300 LAC was grown overnight in RPMI 1640 (Gibco, cat. no. 11875119) at 37°C, slanted with 200 rpm agitation^62^. In both cases, media was supplemented with 1 g/L of ^12^C or U^13^C glucose and sterile filtered with a 0.22-μm filter. After overnight culture, bacteria were heat-killed by incubating at 70°C for 1 h. Heat-killed bacteria were washed 3 times with ultrapure water and aliquots were pelleted and frozen at -80°C to be thawed immediately prior to experiments. Complete heat-killing was confirmed by negative culture after 72 h colony forming unit (CFU) plating on Luria broth agar (*E. coli*) or tryptic soy broth agar (*S. aureus*).

### Lipid and metabolite extraction

At experimental endpoints, media was aspirated from 12-well tissue culture plates, cells were washed twice with ice-cold 0.9% NaCl solution, and metabolites were extracted with 500 μL of 80% methanol on ice. During extraction, 0.5 ng of norvaline and 1 ng of deuterated palmitate (C16:0-d_31_) were added to samples as internal standards for metabolite and lipid recovery, respectively. Wells were scraped, and 10% of each sample was collected and dried for protein quantification by BCA assay. To separate extracts into polar and organic phases, 400 μL of chloroform and 100 μL of water were added to samples. Samples were vortexed and centrifuged at 21,000 g for 5 min at 4°C. The upper aqueous phase was transferred to a gas chromatography-mass spectrometry (GC-MS) vial and dried using a centrivap at 4°C overnight and stored at - 80°C. The lower organic phase was collected and dried under nitrogen gas and stored at -80°C.

### GC-MS analysis of polar metabolites

Dried polar metabolites were processed for GC-MS as previously described^63^. Briefly, polar metabolites derivatization was achieved using a Gerstel MultiPurpose Sampler (MPS 2XL). Keto-containing metabolites were first protected through methoximation by adding 15 μL of 2% (w/v) methoxylamine hydrochloride (MP Biomedicals; cat. no. 02155405-CF) in pyridine and incubating at 45°C for 60 min. Samples were then derivatized by adding 15 μL of N-tert-butyldimethylsily-N-methyltrifluoroacetamide (MTBSTFA) with 1% tert-butyldimethylchlorosilane (tBDMS) (Regis Technologies, Morton Grove, CAS no. IL77377-52-7) and incubating at 45°C for 30 min, generating volatile derivatives suitable for GC-MS analysis. Derivatized polar samples were injected into a GC-MS system equipped with a DB-35MS column (30 m x 0.25 mm x 0.25 μm; Agilent J&W Scientific) installed in an Agilent 7890B GC system integrated with an Agilent 5977A MS. Samples were injected at a GC oven temperature of 100°C which was held for 2 min before ramping to 320°C at 10°C/min and held for 4 min. Electron impact ionization was performed with the MS quadrupole scanning over the range of 100 to 650 mass/charge ratio (m/z) for polar metabolites. Metabolite levels and mass isotopomer distributions were analyzed with an in-house MATLAB script which integrated the metabolite fragment ions and corrected for natural isotope abundances.

### GC-MS analysis of bacterial fatty acid methyl esters (FAMEs)

Dried lipid extracts were prepared from 1e8 CFU equivalents of ^12^C or ^13^C HKEC or HKSA, generated as described above for mammalian cell lipid extraction with C16:0-d_31_ added as an internal standard. Fatty acid methyl esters (FAMEs) were generated as previously described with some modifications^64^. Briefly, lipid extracts were methanolyzed in acidic methanol (4% HCl) for 2 h at 50°C. After derivatization, 1:5 saturated NaCl solution was added to each sample followed by 1:1 hexane. Samples were vortexed and the top organic fraction was collected. This process was repeated once more with fresh hexane and the organic fractions were pooled and dried under nitrogen gas. Just prior to GC-MS analysis, samples were resuspended in 70 μL of hexane and transferred to glass inserts in GC-MS vials. GC-MS analysis of FAMEs was achieved using a Select FAME column (100 m × 0.25 mm × 0.25 µm) installed in an Agilent 7890 A GC interfaced with an Agilent 5975 C MS using the following temperature program: 80°C initial, increase by 20°C/min to 170°C, increase by 1°C/min to 188°C, then 20°C/min to 250°C and hold for 10 min. Mass spectra were acquired using a quadrupole MS with a scan range of m/z 120 to 400. Fatty acid identities were confirmed by comparing the peaks to a standard mixture using retention time and fragmentation patterns. Specifically, cyclopropane fatty acid cyC17:0, cis-vaccenic acid, trans-vaccenic acid, and *anteiso*-C15:0 fatty acids were derivatized alongside bacterial extracts as external standards with the Supelco 37 component FAMEmix analyzed directly. Fatty acid levels and mass isotopomer distributions were analyzed with an in-house MATLAB script which integrated the characteristic lipid fragment ions and corrected for natural isotope abundances.

### Lipid Profiling by LC-MS

Lipid extracts were resuspended in 100 μL of buffer B described below, and 5 μL was injected for analysis. Lipid separation was performed on a Kinetex® C18 column (2.1 × 100 mm, 1.7 μm; Phenomenex) maintained at 35 °C, using a Vanquish HPLC system (Thermo Fisher Scientific). The mobile phases were as follows: buffer A, 2:98 methanol:water with 5 mM ammonium acetate; buffer B, 1:1 methanol:isopropanol with 5 mM ammonium acetate. A typical 50-minute LC run was carried out at a flow rate of 0.2 mL/min. Following injection, the mobile phase was held at 30% buffer B for 1 minute, ramped to 70% B between 1 and 2 minutes, and gradually increased to 95% B from 2 to 13 minutes. The gradient was held at 95% B from 13 to 30 minutes, followed by re-equilibration to 30% B from 30 to 40 minutes. Mass spectrometry was performed on a Thermo Q Exactive Orbitrap (Thermo Fisher Scientific) using heated electrospray ionization (HESI) with polarity switching to acquire full MS1 scans in both positive and negative ion modes within a single LC run. The spray voltage was set to +3.5 kV and –2.5 kV, with a capillary temperature of 325 °C and a probe heater temperature of 400 °C. Full MS scans were acquired at a resolution of 70,000 (at m/z 200), with an AGC target of 1 × 10⁶, maximum injection time of 150 ms, and a scan range of m/z 150–1500. A representative subset of samples was analyzed using data-dependent MS² (dd-MS²) acquisition in dedicated positive and negative ion mode runs performed separately on a pooled sample. MS² scans were acquired at a resolution of 17,500 with an AGC target of 1 × 10⁵, maximum injection time of 50 ms, isolation window of 1.2 m/z, and normalized collision energy of 30. The top 12 precursor ions were selected for fragmentation, with dynamic exclusion set to 10 s and isotope exclusion enabled. Extracted ion chromatograms (EICs) were generated in El-MAVEN ±5 ppm mass tolerance.

### Lipid isotopologue analysis

Lipid LC-MS data were analyzed using EI-Maven. Mass error was set to ±5 ppm for lipid identification. Mass isotopologues were filtered to elute within 5 seconds of the unlabeled parent ion. Only MS^2^ confirmed lipids were included in analysis, and identification was based on lipid species-specific fragmentation patterns predicted in LipidCreator and confirmed with external standards, including UltimateSPLASH™ ONE (Avanti Polar Lipids, cat. no. A83820), EquiSPLASH™ (Avanti Polar Lipids, cat. no. A83731), and an acylcarnitine standard mix (Cambridge Isotope Laboratories, cat. no. NSK-B-US-1). Relative abundance was calculated by normalizing to internal standard (C16:0-d_31_) and protein levels then dividing by the average of the control condition specified in figure legends. Correction for the natural abundance of ^13^C was performed with an in-house MATLAB script. For most analysis, the relative abundance of selected, ^13^C-corrected isotopologues corresponding to recycling of intact bacteria-derived fatty acids into the macrophage lipidome are reported without further processing. For lipidomic analysis, peaks were filtered for a minimal intensity of 1e5 ion counts and a signal-to-noise ratio >10. To reduce matrix-associated noise in ^13^C bacterial free fatty acid quantification and full lipid mass isotopologue distributions of ^13^C bacteria-derived carbon salvage into the macrophage lipidome (**Fig S1**), paired unlabeled macrophage spectra were treated as matrix blanks to subtract from labeled sample spectra post-isotope correction.

### Polar metabolite profiling by LC-MS

Chromatographic separation was achieved using a Vanquish Flex UHPLC (Thermo Fisher Scientific) system equipped with a InfinityLab Poroshell 120 HILIC-z column (2.1 ×100 mm, 2.7 µm; Agilent). Mobile phases A and B were comprised of 10 mM ammonium carbonate in water (A) and in 95:5 acetonitrile: water (B) with 5µM medronic acid. A flow rate of 0.3 mL/min was used to elute compounds with the following gradient: 100% B, decreasing to 90% B over 4 min, to 50% B over 6 min, then to 30% B in 0.5 min, held for 1 min before returning to 100% B and equilibrating for 8 min. The column was maintained at 45°C. The heated electrospray ionization (HESI) conditions used were as follows; spray voltage, 3.9 kV (ESI+), 3.5 kV (ESI−); capillary temperature, 250 °C; probe heater temperature, 350 °C; sheath gas, 46 arbitrary units; auxiliary gas, 11 arbitrary units; and S-Lens RF level, 50%. Full MS data were collected using a Q-Exactive Quadrupole Orbitrap mass spectrometer (Thermo Fisher Scientific) in polarity switching ionization mode separately from mass ranges 70–1000 m/z for both negative and positive mode, at 70,000 resolution. The automatic gain control (AGC) was set to 1 ×106 and maximum injection time (IT) used was 150 ms. Top 12 data dependent MS/MS (ddMS2) spectra were also collected for a representative pool of unlabeled samples with MS1 and MS2 data acquired at 35,000 and 17,500 resolution, respectively. The AGC target and maximum IT in the ddMS2 experiment were set to 5×105, 100 ms for MS1 spectra and 1×105, 50 ms for the MS2 spectra. A 1.2 Da isolation window and normalized collision energy of 25 were used for ddMS2 with the underfill ratio set to 1.0% along with 10 sec dynamic exclusion to reduce redundant spectra.

### Immunofluorescence assay

For immunofluorescence assays (IFA), BMDMs or peritoneal exudate cells were seeded overnight in optical bottom imaging-grade 96-well plates (PhenoPlate; Revvity Health Sciences, cat. no. 6055302). Cells were fixed with 4% paraformaldehyde (PFA) at room temperature (RT) for 15 min and permeabilized with PBS containing 0.1% Triton-X100 for 20 min at RT. Cells were blocked with blocking buffer for 30 min at RT with (PBS containing 5% BSA (Sigma-Aldrich, cat. no. A9647) and 10% normal goat serum (Invitrogen, cat. no. 10000C). Cells were incubated with primary antibodies (1:1000) in blocking buffer for 1 h at RT. Cells were washed with PBS and stained for 30 min with secondary antibodies (1:1000) in blocking buffer with Hoechst 33342 (10 μg/mL; Invitrogen, cat. no. H3570) or DAPI (1 μg/mL; Invitrogen, cat. no. D1306) as a nuclear counterstain. For combined C12-BODIPY tracing and IFA, live cells were incubated with HKEC particles (MOI 25) loaded with C12-BODIPY generated as described below. At the endpoint, C12-BODIPY HKEC-fed cells were fixed with 4% PFA at RT for 15 min and then permeabilized with 20 μM digitonin in KHM buffer (110 mM potassium acetate, 20 mM HEPES, 2mM MgCl2) for 2 min then washed with PBS. IFA and imaging were performed on the same day as the experimental endpoint. Primary antibodies used for IFA in this study were rabbit IgG anti-*E.* Coli (Bio-Rad, cat. no. 4329-4906), mouse IgG anti-RAB7 (Cell Signaling Technology, cat. no. E907E), and rabbit IgG anti-TOMM20 (Abcam, cat. no. ab186734). Secondary antibodies used for IFA in this study were AlexaFluor 488 (AF488)-conjugated goat anti-rabbit IgG (Thermo Fisher Scientific, cat. no. A11008) and AF647-conjugated goat anti-mouse IgG (Thermo Fisher Scientific, cat. no. A21235).

### Generation of C12-BODIPY-loaded heat-killed bacterial particles

HKEC were generated as described above for the ^12^C HKEC control in stable isotope tracing experiments. Frozen aliquots of HKEC were freshly thawed before each experiment. Zombie Violet (Biolegend, 423113) dye was prepared by dissolving lyophilized powder in 100 μL DMSO and then incubated with bacterial particles at a 1:100 dilution for 30 min at RT. At the endpoint, staining was quenched with DMEM + 10% FBS then washed with ultrapure water. Zombie Violet-labeled HKEC were incubated with 100 μg/mL C12-BODIPY (Cayman Chemical, cat. no. 27014) at RT for 10 min and washed with ultrapure water. Unless specified, the Zombie Violet conjugation step was omitted from C12-BODIPY HKEC generation.

### Automated high-content confocal microscopy

Automated high-content spinning disk confocal microscopy was performed using a CQ3000 high-content microscope (Yokogawa) or a BioTek Cytation C10 confocal imaging reader (Agilent). On the Cytation C10, imaging was performed with an Olympus 60x air interface objective and a Hamamatsu ORCA CMOS digital camera. Single optical sections were imaged for each acquisition field with laser-defined autofocus. Nine acquisition fields were randomly selected per well. Representative images have been deconvolved using 5-iteration default deconvolution in Gen5 software. On the CQ3000, imaging was performed with a 60x/1.2NA objective with water immersion and dual sCMOS cameras. Maximum intensity projection (MIP) images were collected across 10-15 z-planes spanning 6-10 μm depending on sample depth and centered around a laser autofocus-defined focal plane for each acquisition field. For each experimental condition, 6-15 fields of view per well were randomly selected with the goal to achieve >100 cells sampled per biological replicate. Representative MIPs have been processed with 50-pixel rolling ball radius background subtraction and 1-pixel gaussian blur filtering in ImageJ. Representative images within each experiment have been identically processed and uniformly scaled in brightness and contrast between groups.

### Quantitative microscopy analysis

Open-source image analysis software Cellprofiler was used for automated analysis of all micrographs generated. Cellprofiler pipelines can be found in the supplemental files (**File S1-S4**). Image analysis was performed on MIP images generated in CellPathfinder software or raw confocal images from Gen5 software. Automated single-cell analysis was achieved by segmentation of nuclear objects based on the intensity of the nuclear stain (DAPI or Hoechst) using global two-class Otsu thresholding in the identify primary objects module, followed by propagation of the nuclear objects to the cellular periphery based on a smoothing of the mitochondrial or endolysosomal stain to give a rough cytoplasmic area. In live imaging experiments analyzing the extraction of C12-BODIPY from Zombie Violet-stained HKEC, brightfield images were inverted, smoothed, then segmented with global two-class Otsu thresholding in the identify primary objects module to define single cells. In phagocytosis experiments, the anti-*E. coli* immunostain or the Zombie Violet stain in labeled HKEC was segmented using the identify primary objects module with two-class Otsu-based adaptive thresholding, and the bacteria-positive area was summed and related to parent cells using the relate objects module. To achieve lipid droplet analysis, focal regions of BODIPY493 intensity were isolated from the background neutral lipid stain based on the enhance features module with speckles enhancement with 12-pixel feature size selection. Subsequently, isolated lipid droplet foci were segmented using the identify primary objects module with two-class Otsu-based adaptive thresholding. Following cellular and lipid droplet segmentation, the number of lipid droplets and the cumulative intensity of BODIPY493 and C12-BODIPY within each lipid droplet were measured and related to the parent cells using the relate objects module. Colocalization of C12-BODIPY with mitochondrial stains, MitoTracker Deep Red (Invitrogen, cat. no. M22426) or anti-TOMM20 (Abcam, cat. no. ab186734) was performed on a single cell basis using the measure colocalization module with Pearson’s correlation reported.

### *Ex vivo* challenge of *in vivo* trained peritoneal macrophages

For *ex vivo* peritoneal macrophage experiments, WT mice were challenged with intraperitoneal injection (IP) of LPS (2 mg/kg O111:B4) or PBS control and recovered for 1, 3, 5, or 7 days. At endpoint, mice were euthanized and peripheral blood was collected for analysis of cytokines (as described below). Cells were isolated by washing the peritoneal cavity with 3 mL of PBS. Peritoneal exudates were centrifuged, resuspended in D10, and seeded into 12-well tissue culture plates or 96-well imaging grade plates. Cells were allowed to adhere for 1 h, then non-adherent cells were washed away with PBS. Remaining cells were verified to be macrophages using immunofluorescence staining of F4/80 in paired plates that were imaged with a CQ3000 high-content confocal microscope. Post-wash, peritoneal macrophages were stimulated with either ^13^C or C12-BODIPY-loaded HKEC for 24 h in D10. At the endpoint, C12-BODIPY HKEC-fed peritoneal macrophages were processed according to the immunofluorescence and live imaging protocols (outlined above), and ^13^C HKEC-fed macrophages were extracted (metabolite extraction protocol outlined above).

### *In vivo* training and challenge with ^13^C heat-killed *E. coli*

WT mice were challenged with IP injection of LPS (2 mg/kg O111:B4) or PBS control. Day 5 post-injection was selected for recovery because it showed the most pronounced *ex vivo* lipid catabolic phenotype among peritoneal macrophages. After recovery, mice were challenged for 24 h with 1 x 10^10^ of ^13^C HKEC, a dose that was determined in a pilot study to be the LD_30_ in naïve animals. Clinical features were monitored longitudinally including weight, rectal temperature, and clinical score, using the scoring system defined below: 5. Normal exploratory behavior, rearing on hind limbs, and grooming; 4. Reduced exploratory behavior, rearing on hind limbs, and grooming. Slower and/or less steady gait, but free ambulation throughout the cage; 3. Slow, unsteady gait got <5 s; 2. No voluntary movement, but the mouse can generate slow, unsteady, gait for > 5 s; 1. The mouse does not move away from stimulation by research, but can still right itself; 0. Deceased. Rectal temperatures were binned based on the following strategy (below) and summed with morbidity scores (above) to calculate overall health scores at each timepoint for health trajectory plots (5. 35->38°C, 4. 31-34.9°C, 3. 28-30.9°C, 2. 25-27.9°C, 1. 22-24.9°C, 0. <22°C). At the endpoint, the peritoneal exudate was collected, the cell pellet was washed, and cellular lipids were extracted and analyzed with LC-MS.

### ELISA and Luminex analysis of secreted and peripheral cytokines

Supernatants from naïve or LPS-trained BMDMs stimulated ± HKEC were collected and clarified by centrifugation at 1000 x g for 5 min at 4 °C. The clarified supernatant was immediately stored at -80 °C. *In vivo* peripheral cytokine production post-LPS or PBS control recovery was measured from serum isolated from cardiac puncture at day 1, 3, 5, and 7. *In vitro* and *in vivo* samples were shipped to Eve Technologies (Calgary, Alberta, Canada) for luminex-based cytokine profiling. Cytokines with values below 20 pg/mL were considered below the limit of detection and filtered. *In vitro* cytokine measurements are presented as the Log_2_ fold-change compared to the average of the naïve + HKEC condition. *In vivo* cytokine measurements are presented as the mean fold change compared to D1 post-LPS cytokine abundance. Measurements of targeted cytokines TNF and IL-10 were achieved by commercially available enzyme-linked immunosorbent assays (ELISA) analysis (Proteintech, cat. no. KE10002 and KE10103, respectively) of clarified supernatants from WT and IL-10 KO naïve and LPS-trained BMDMs.

### SDS-PAGE and Immunoblot

BMDMs were washed with ice-cold PBS and then lysed with RIPA lysis buffer supplemented with a protease/phosphatase inhibitor cocktail (Cell Signaling Technology, cat. no. 9806 and 5872, respectively) for 15 min on ice. Lysates were centrifuged at 15,000 g for 20 min at 4°C. Following extraction, sample protein content was measured by BCA assay with bovine serum albumin as standard (Bio-Rad), and protein content was normalized by dilution. Subsequently, samples were diluted in Laemmli sample loading buffer, heated for 5 min at 95°C, and then separated on 4-20% gradient polyacrylamide tris-glycine gels. After SDS-PAGE, protein was transferred to a methanol-activated polyvinylidene fluoride (PVDF) membrane with a wet transfer system using tris-glycine transfer buffer for 1 h at 100 V at 4 °C. Membranes were blocked with 5% BSA in Tris-buffered saline (TBS) with 0.1% Tween 20 (IB blocking buffer) for 30 min at RT and then incubated with primary antibodies in blocking buffer overnight at 4°C. Blots were washed 3 times in TBS with 0.1% Tween 20, incubated with anti-rabbit IgG horseradish peroxidase (HRP)-conjugated (Cell Signaling Technology, cat. no. 7074) or anti-mouse StarBright700-conjugated secondary antibodies (1:10,000; Bio-Rad, cat. no. 12004158) for 1 h at RT, washed, and then developed with Prosignal Femto ECL reagent (Genesee Scientific, cat. no. 20-302B) or fluorescence imaging, respectively, on a Chemidoc MP Imaging System (Bio-Rad). Primary antibodies used for immunoblot in this study include anti-ACOD1 rabbit IgG (Abcam, cat. no. ab222411) and anti-ACTB Mouse IgG (Cell Signaling Technology, cat. no. 3700). Immunoblots were quantified using the ImageJ densitometric gel analysis protocol for 1D gels.

### Seahorse extracellular flux assay

The oxygen consumption rate (OCR) of BMDMs was measured using an Agilent Seahorse XF96 analyzer. BMDMs were plated in Seahorse 96-well assay plates and cultured overnight in DMEM supplemented with 10 mM glucose, 1 mM pyruvate, 2 mM glutamine, and 10% FBS. A target cell density of 80% confluence was achieved by addition of 45,000 BMDMs per well. After the LPS training described above, medium was exchanged for Seahorse DMEM supplemented with equivalent levels of glucose, pyruvate, and glutamine without FBS. Cells were kept in a 37°C incubator without CO_2_ for 30 min before analysis. In all experiments, the Mito Stress Test assay was used with 1.5 μM oligomycin (O), 5 μM carbonyl cyanide m-chlorophenyl hydrazone (C), 0.5 μM rotenone (R), and 0.5 μM antimycin A (AA). In a subset of experiments, cells were pre-treated with ETO (10 μM) for 4 h prior to analysis. OCR measurements are normalized to well-paired protein measurements determined by BCA assay.

### RNA sequencing

Naïve and LPS-trained WT and IL-10 KO BMDMs were prepared according to the *in vitro* training protocol (outlined above). At endpoint, cells were washed with PBS and resuspended in 50 μL DNA/RNA Shield (Zymo Research Corp, cat. no. R1100-250) and submitted for RNA sequencing analysis using Plasmidsaurus’ 3’-end counting RNA sequencing service (Louisville, KY, USA)..

Briefly, poly(A)+ mRNA was captured using an anchored poly(dT)VN primer containing unique molecular identifiers (UMIs). Subsequently, cDNA was generated by reverse transcription and second strand synthesis, followed by tagmentation, library indexing, and PCR amplification introducing unique dual indices (UDIs). Resulting libraries were single-end sequenced on an Illumina NovaSeq X Plus instrument using a stranded 3’-end counting method. Sequencing reads were processed with the Plasmidsaurus bioinformatics pipeline. Raw sequencing data was demultiplexed, quality filtered, aligned to the GRCm39 mouse reference genome with STAR aligner v2.7, and coordinate-sorted. Sequencing data was deduplicated by UMI grouping. Gene-level expression was quantified with featureCounts (Subread package v2.1.1) with strand-specific counting, fractional assignment of multi-mapping reads, and exon and 3’ untranslated region feature grouping. Gene-level count tables were generated and annotated, and differentially expressed genes (DEGs) were identified using edgePython v0.2.5. Mitochondria-associated DEGs were filtered using the MitoCarta3.0 gene list^65^.

### Statistical analysis

GraphPad Prism was used for the majority of statistical analysis, with individual tests indicated in figure legends. In general, pair-wise comparisons were analyzed using an unpaired *t*-test, multiple comparisons were analyzed using one-way ANOVA with Sidak’s post-test, and grouped analysis was performed using two-way ANOVA with Tukey’s post-test. The *p*-values displayed are adjusted for multiple comparisons. In cases where experiment-level replicates are shown, individual data points represent distinct biological sources (e.g. different mice).

## Extended Data Figures

**Fig S1.**
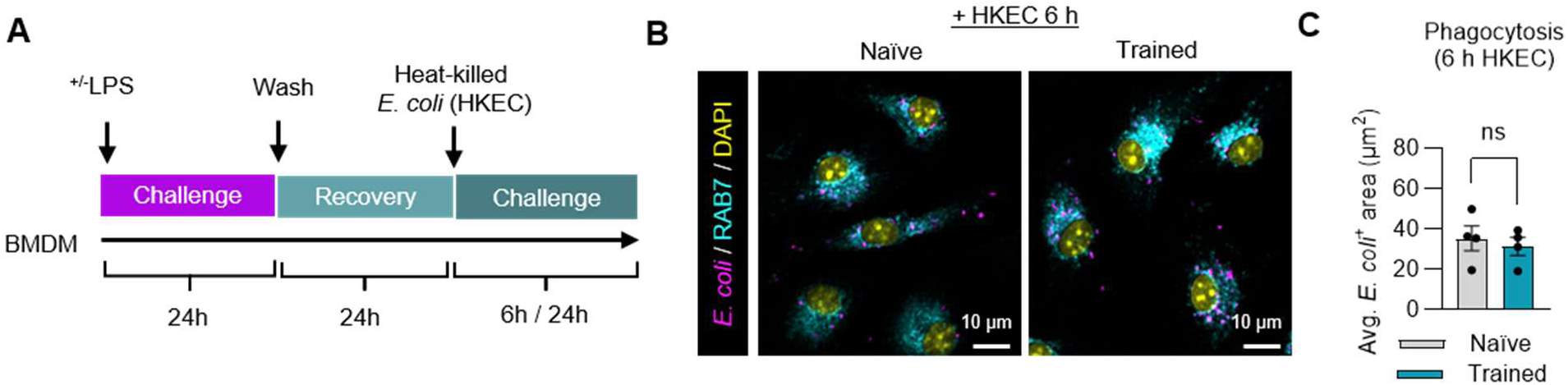
Effect of TLR4 training on heat-killed *E. coli* phagocytosis. **A.** Schematic representation of *in vitro* Toll-like receptor 4 (TLR4) training of murine bone marrow-derived macrophages (BMDMs), which follows a 24 h primary stimulation with (trained) or without (naïve) 100 ng/mL O111:B4 lipopolysaccharide (LPS), a 24 h recovery period, then a re-challenge period between 6 to 24 h with heat-killed wild-type K12 MG1655 *E. coli* (HKEC) at multiplicity of 25 heat-killed particles per host cell. Unless otherwise indicated, this training and challenge model is unmodified throughout subsequent experiments. **B.** Representative confocal micrographs from immunofluorescence labeling of the late endolysosomal marker RAB7, *E coli* antigen, and a nuclear counterstain (DAPI) in naïve and trained BMDMs fed HKEC for 6 h. **C.** High-content automated quantitation of average bacterial load per cell calculated as the area within each cell occupied by segmented *E. coli* antigen signal. *p*-values were calculated with an unpaired *t*-test. ns > 0.05.

**Fig S2.**
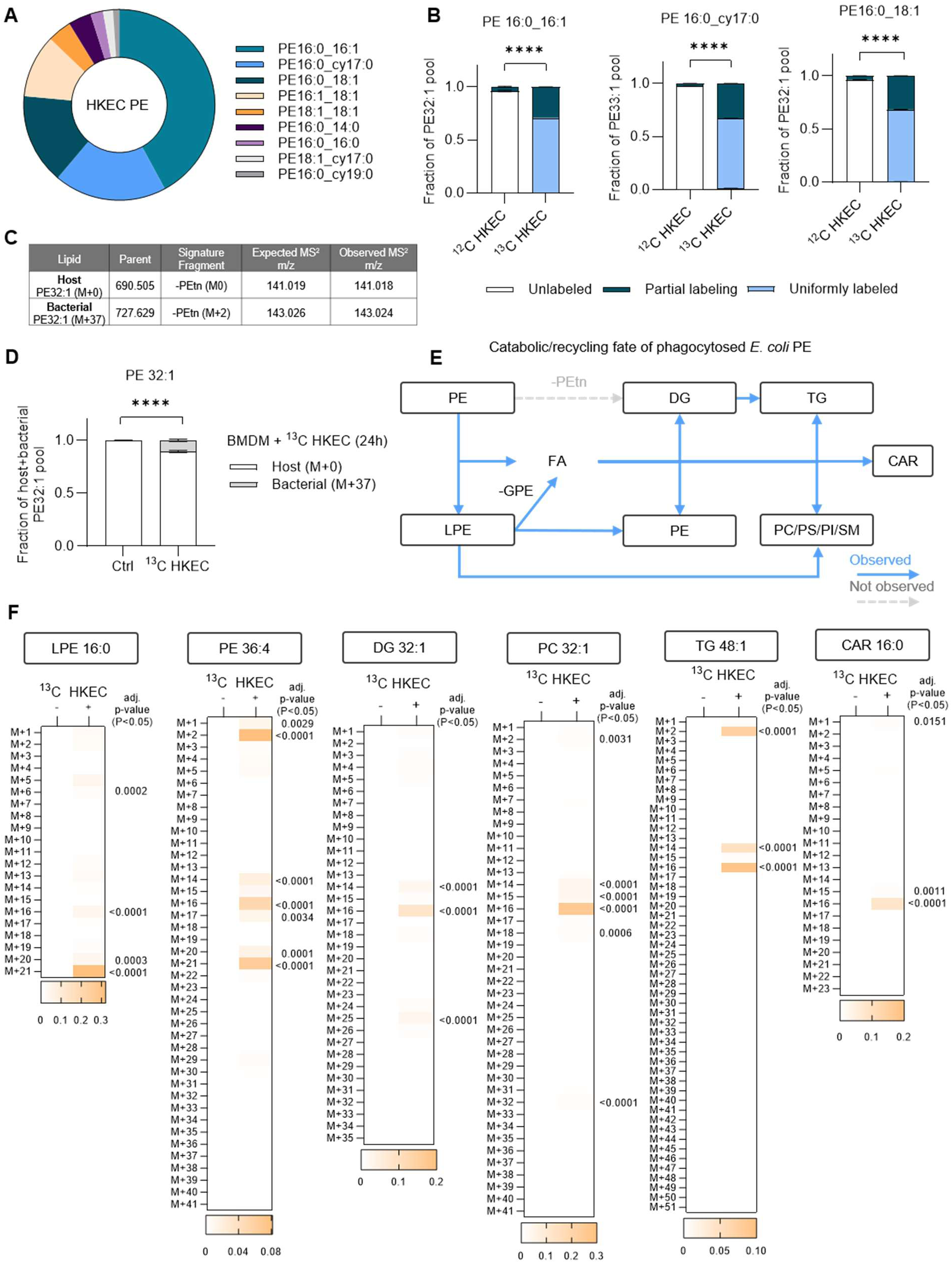
Generation and analysis of ^13^C stable isotope labeled heat-killed *E coli*. **A.** LC-MS analysis of molar ratios of major phosphatidylethanolamine (PE) species from heat-killed *E coli*. PE fatty acid composition was determined by negative mode MS^2^. **B.** Fractional enrichment of ^13^C in a subset of major *E. coli* PE species in lipid extracts of ^12^C vs ^13^C HKEC. Partial labeling is almost exclusively one less ^13^C than U^13^C. **C.** In a BMDM-HKEC co-incubation system, host and U^13^C *E. coli*-derived PE can be distinguished based on parent mass and positive mode MS^2^ neutral loss of an M+2 phosphoethanolamine in ^13^C HKEC condition. **D.** Relative proportion of host-derived and bacteria-derived PE32:1 following incubation of WT naïve BMDMs with ^13^C HKEC for 24 hours. **E.** Bacteria-derived PE catabolic and recycling pathways observed in macrophages, with blue arrows indicating pathways that are supported by stable isotope tracing results and the dashed grey arrow indicating a reaction that is not detected in this system. Abbreviations: diacylglycerol (DG), triacylglycerol (TG), phosphatidylcholine (PC), phosphatidylethanolamine (PE), lysophosphatidylethanolamine (LPE), phosphatidylserine (PS), phosphatidylinositol (PI), and sphingomyelin (SM), acylcarnitine (CAR), phosphoethanolamine (Petn), glycerophosphoethanolamine (GPE), and fatty acid (FA). **F.** Mass isotopologue distributions for LPE 16:0, PE 36:4, DG 32:1, PC 32:1, TG 48:1, and CAR 16:0 from WT naïve BMDMs treated for 24 h ± ^13^C HKEC. LPE 16:0 M+21 reflects a U^13^C species directly derived from a bacterial PE through lipase activity. M+5 in the LPE pool reflects GPE recycling. M+16 in the LPE and all other lipid pools reflects C16 fatty acid recycling from bacterial lipids. M+21 in the PE36:4 pool reflects reacylation of a LPE with a host-derived FA. Finally, M+2 reflects elongation of host fatty acids with bacteria-derived acetyl-CoA. While theoretically M+2 could represent recycling of ethanolamine into the PE headgroup, MS^2^ analysis of Petn neutral loss from M+2 PE species does not carry a label, indicating that this type of recycling is uncommon in this system. For **A-D**, graphs represent the mean of 3 technical replicates used to validate HKEC frozen stocks. For **F,** graphs represent the mean of 11 biological replicates. *p*-values were calculated using two-way ANOVA with Sidak’s post-test (**B**, **D**, **F**). ns > 0.05; ∗*p* < 0.05; ∗∗*p* < 0.01; ∗∗∗*p* < 0.001; ∗∗∗∗*p* < 0.0001.

**Fig S3.**
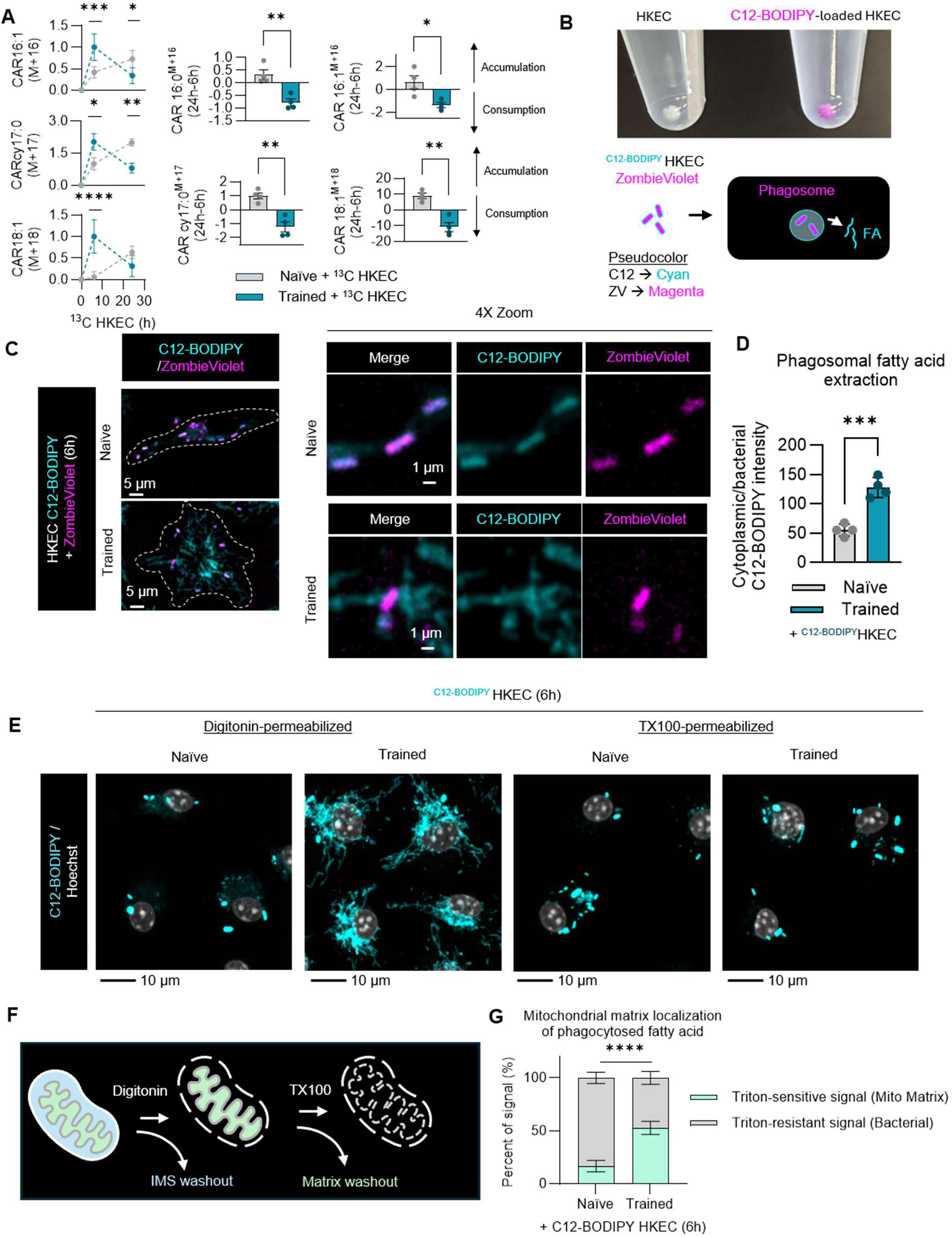
Generation and analysis of C12-BODIPY fluorescent fatty acid-labeled heat-killed *E coli*. **A.** LC-MS analysis of bacterial fatty acid (C16:1, C18:1, cyclopropane C17:0) catabolism via the carnitine shuttle, measured as either kinetic analysis of the relative abundance of ^13^C bacterial fatty acid-loaded acylcarnitine (CAR) or the ratio of bacterial fatty-loaded CAR at 24 h relative to the abundance at 6 h post-stimulation. **B.** Macroscopic photograph of control HKEC and C12-BODIPY-loaded HKEC pellets in PBS and schematic representation of general bacterial particle labeling with amine cross-linked fluorophore Zombie Violet and specific fatty acid labeling with C12-BODIPY. Pseudocoloring of C12-BODIPY to cyan is maintained throughout the manuscript. **C.** Representative maximum intensity projection of Z-stack confocal micrographs of naïve and trained BMDMs fed with Zombieviolet and C12-BODIPY-labeled HKEC for 6h and analyzed by live cell imaging. Cell outlines were drawn based on paired phase contrast images. Zoomed regions of interest demonstrate retention of C12-BODIPY within bacterial particles in naïve cells and extraction of C12-BODIPY into the cytoplasm of TLR4-trained cells**. D.** Quantification of the average ratio of cytoplasmic:bacterial C12-BODIPY per cell. **E.** Representative maximum intensity projection of Z-stack confocal micrographs of C12-BODIPY and Hoechst nuclear counterstain from naïve and trained BMDMs fed C12-BODIPY-loaded HKEC for 6 hours followed by a differential strength detergent permeabilization assay, comparing digitonin (plasma membrane and outer mitochondrial membrane permeabilization) and Triton X-100 (total membrane permeabilization). **F.** Cartoon diagram demonstrating compartmentalization interpretations for differential detergent assay. **G.** Calculation of the Triton-sensitive (mitochondrial matrix) and Triton-resistant (bacterial particle-associated) fractions of total fluorescence in naïve and trained BMDMs fed C12-BODIPY-loaded HKEC for 6 h. Graphs represent the mean of n = 4 biological replicates with SEM error bars for all graphs. *p*-values were calculated using an unpaired *t*-test (**A, D**) and two-way ANOVA with Sidak’s post-test (**A**, **G**). ns > 0.05; ∗*p* < 0.05; ∗∗*p* < 0.01; ∗∗∗*p* < 0.001; ∗∗∗∗*p* < 0.0001.

**Fig S4.**
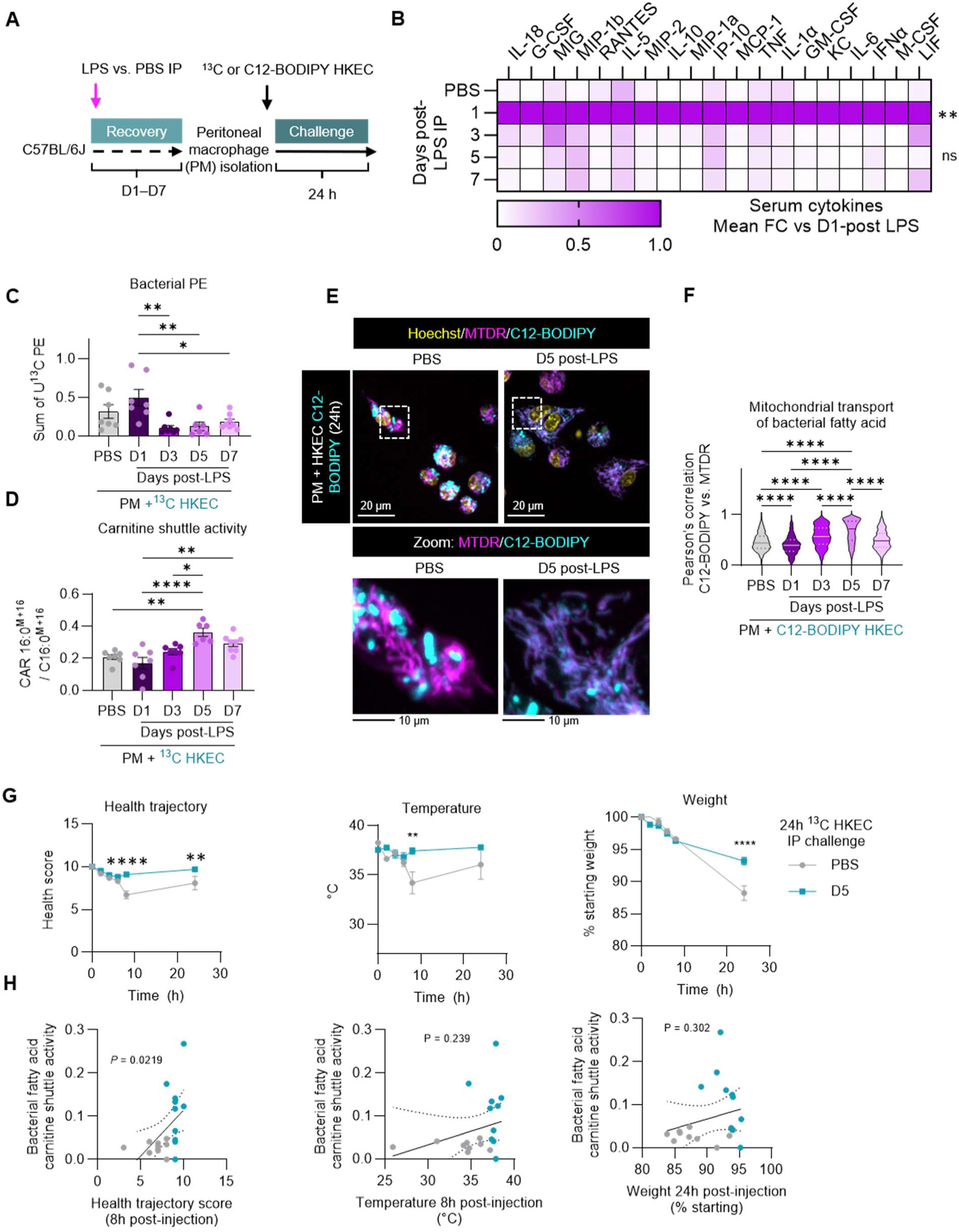
Recovery from sublethal endotoxemia improves macrophage catabolism of bacterial lipids. **A.** Experimental design of *ex vivo* assessment of peritoneal macrophage (PM) lipid catabolic function post-sublethal intraperitoneal (IP) injection endotoxemia (O111:B4 LPS, 2mg/kg) recovered for 1, 3, 5, or 7 D compared to PBS control. **B.** Luminex analysis of serum cytokines at D1, D3, D5, and D7 post-LPS IP compared to PBS control. Data are presented as the mean fold change over the maximal production which was detected at D1 for all cytokines. LC-MS analysis of summed bacteria-derived phosphatidylethanolamine (U^13^C PE) (**C**) and the ratio of bacterial fatty acid-loaded palmitoylcarnitine (M+16) to bacteria-derived free palmitate (M+16) (**D**) from isolated PMs. **E.** Representative maximum intensity projection of Z-stack confocal micrographs of PMs derived from PBS-injected or D5-post LPS recovered mice fed with C12-BODIPY-loaded HKEC for 24 h and analyzed by live cell imaging with MitoTracker Deep Red (MTDR) and Hoechst counterstains with zoomed region highlighting C12-BODIPY subcellular localization. **F.** Pearson’s correlation coefficient between C12-BODIPY and MTDR, calculated per cell. **G.** Longitudinal measurement of health parameters in PBS or LPS-recovered (D5) mice challenged with 10^10^ HKEC IP for 24 h including temperature, weight and an aggregate score termed overall health score which reflects morbidity, temperature, and weight measurements. **H.** Correlation between health parameters from **G** at indicated time points with acylcarnitine shuttle utilization of bacterial fatty acids. Graphs represent the mean of n ≥ 6 biological replicates with SEM error bars for all graphs except **F**, which represents >650 cells analyzed per condition pooled across 4 biological replicates with violin plots of medians and quartiles. *p*-values were calculated using one-way ANOVA with Tukey’s post-test (**C**, **D**, **F**), two-way ANOVA with Sidak’s post-test (**B**, **G**) and linear regression (**H**). *p*-value designation for **B** denotes the maximum *p*-value across all columns compared to PBS control. ns > 0.05; ∗*p* < 0.05; ∗∗*p* < 0.01; ∗∗∗*p* < 0.001; ∗∗∗∗*p* < 0.0001.

**Fig S5.**
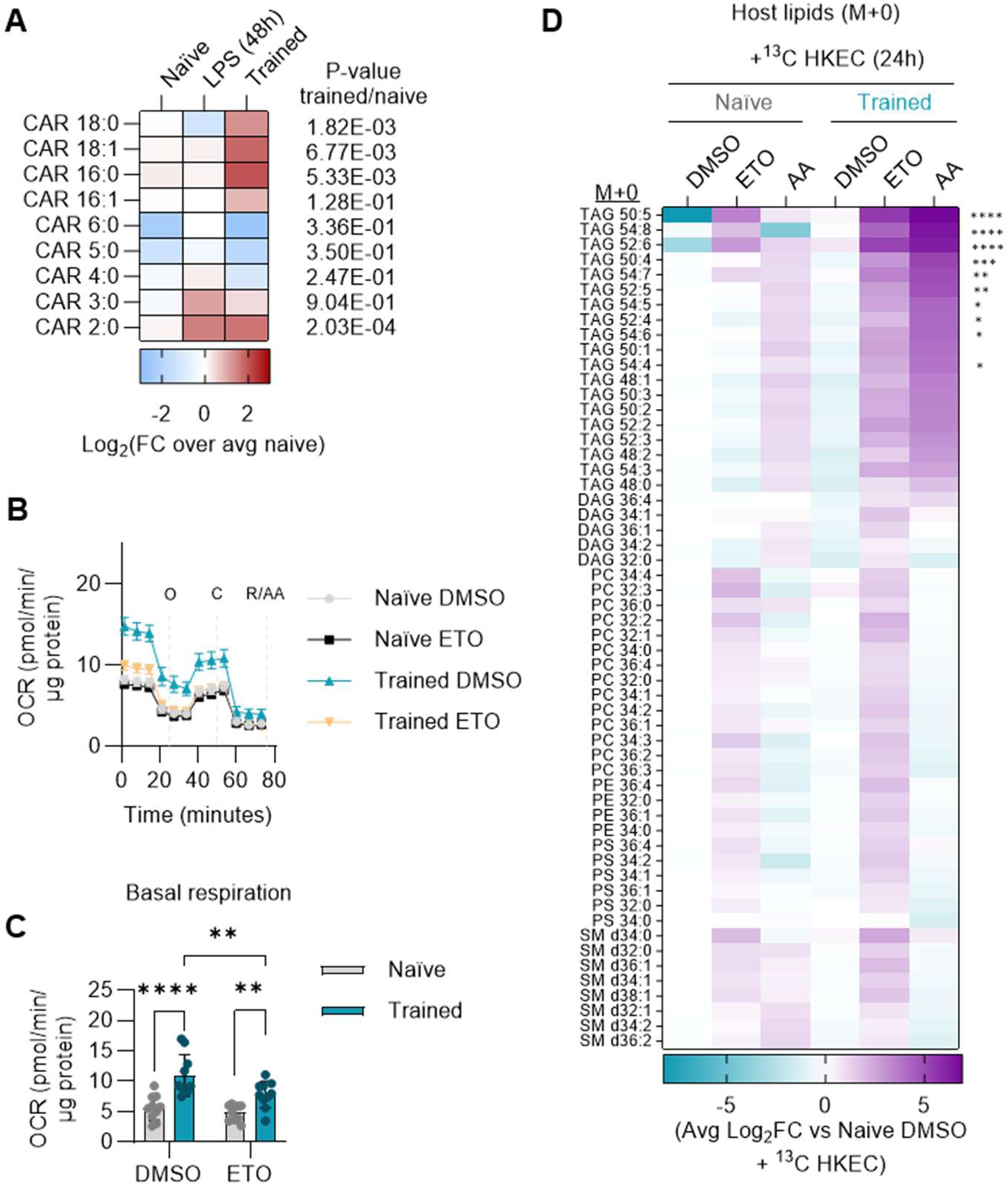
Assessment of carnitine shuttle activity in trained macrophages. **A.** LC-MS analysis of total acylcarnitine pools in naïve, 48 h LPS-stimulated, or TLR4-trained (24 h LPS stimulation followed by 24 h recovery) BMDMs. **B.** Seahorse XF analysis of protein-normalized oxygen consumption rate (OCR) in naïve and trained BMDMs treated with Etomoxir (10 μM) for 4 h using the Mito Stress Test (MST) assay, with additions of oligomycin (O), Carbonyl cyanide m-chlorophenyl hydrazone (C), rotenone (R), and antimycin A (AA) at indicated time points. **C.** Quantification of basal respiration from panel **B. D.** LC-MS analysis of relative abundances of indicated host lipids (M0), corresponding to lipid species highlighted in **Fig 2D** compared between naïve or trained BMDMs stimulated with ^13^C HKEC for 24 h in the presence of ETO, AA, or DMSO and measured as the Log_2_ fold change over naïve BMDMs treated with vehicle control and ^13^C HKEC of relative abundance of individual M0 isotopologues of MS^2^ verified lipids within each lipid class: diacylglycerol (DG), triacylglycerol (TG), phosphatidylcholine (PC), phosphatidylethanolamine (PE), phosphatidylserine (PS), and sphingomyelin (SM). Graphs represent the mean of n ≥ 4 biological replicates with SEM error bars for all graphs except **B**, which represents >35 technical replicate wells analyzed per condition pooled across 10 biological replicates. *p*-values were calculated with a one-way ANOVA with Tukey’s post-test (**A**) and two-way ANOVA with Sidak’s post-test (**B-D**). ns > 0.05; ∗*p* < 0.05; ∗∗*p* < 0.01; ∗∗∗*p* < 0.001; ∗∗∗∗*p* < 0.0001.

**Fig S6.**
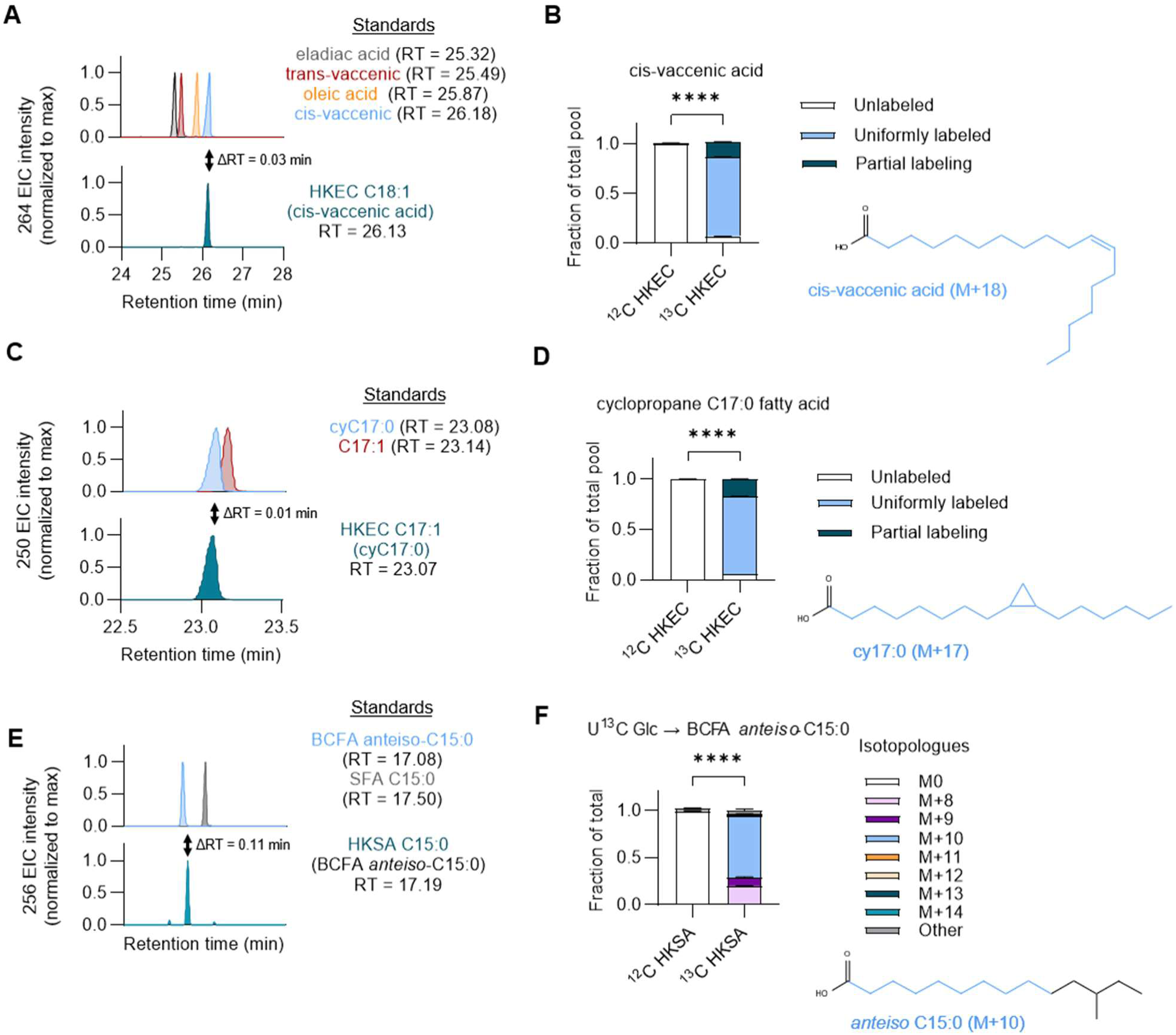
Validation of bacterial fatty acid identity and stable isotope labeling efficiency. **A.** GC-MS analysis of fatty acid methyl ester (FAME)-derivatized lipid extracts from heat-killed *E. coli* (HKEC) alongside derivatized external standards of oleic acid, eladiac acid, cis-vaccenic acid, and trans-vaccenic acid. Extracted ion chromatograms (EIC) from the C18:1 characteristic fragment m/z = 264 is shown for each standard with the closest match to the HKEC sample being cis-vaccenic acid. **B.** Condensed mass isotopologue distribution (MID) from ^12^C vs. ^13^C HKEC of cis-vaccenic acid showing the fraction of the fatty acid pool consisting of unlabeled (M0), uniformly labeled (M+18), and partially labeled (18>M>0) isotopologues. **C.** GC-MS analysis of HKEC FAMEs alongside external derivatized standards of cyclopropane C17:0 (cyC17:0) and cis-10-heptadecenoic acid. EIC from the C17:1 characteristic fragment m/z = 250 is shown for each standard with the closest match to the HKEC sample being cyC17:0. **D.** Condensed MID from ^12^C vs. ^13^C HKEC of cyC17:0 showing the fraction of the fatty acid pool consisting of unlabeled (M0), uniformly labeled (M+17), and partially labeled (17>M>0) isotopologues. **E.** GC-MS analysis of fatty acid methyl ester (FAME)-derivatized lipid extracts from heat-killed *S. aureus* (HKSA) alongside derivatized external standards of pentadecanoic acid (C15:0 straight chain fatty acid [SFA]) and *anteiso*-C15:0 branched chain fatty acid (BCFA). Extracted ion chromatograms (EIC) from the C15:0 characteristic fragment m/z = 256 is shown for each standard with the closest match to the HKSA sample being *anteiso*-C15:0 BCFA. **F.** Condensed MID from ^12^C vs ^13^C glucose-traced HKSA of *anteiso*-C15:0 BCFA showing isotopologues. Graphs represent the mean of 3 technical replicates used to validate HKEC and HKSA frozen stocks with SEM error bars. *p*-values were calculated using a two-way ANOVA with Sidak’s post-test (**B**, **D**, **F**). ns > 0.05; ∗*p* < 0.05; ∗∗*p* < 0.01; ∗∗∗*p* < 0.001; ∗∗∗∗*p* < 0.0001.

**Fig S7.**
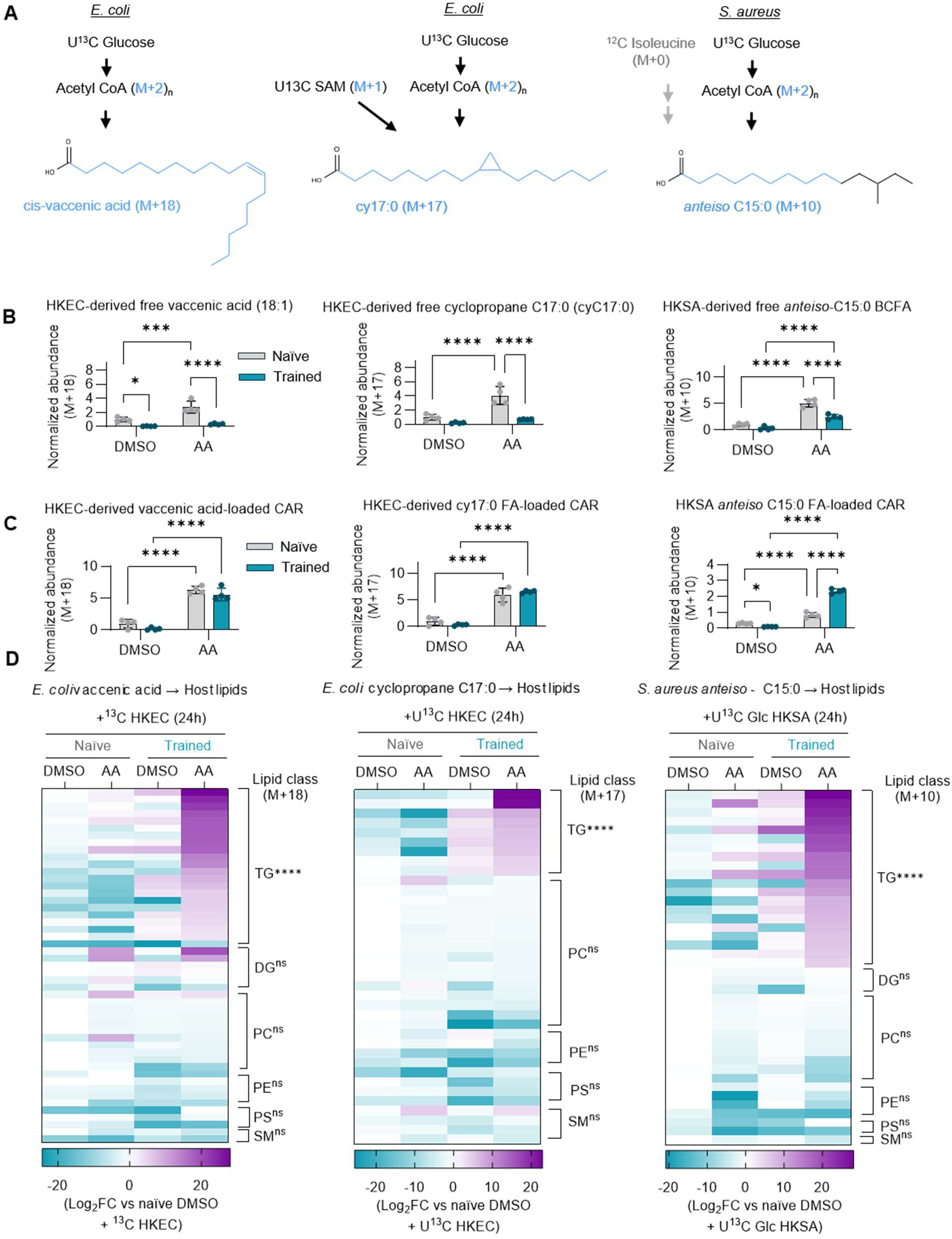
Incorporation of bacterial fatty acids into the host lipidome post-OXPHOS blockade in trained macrophages. Structural illustration and observed isotopic enrichment of analyzed bacterial fatty acids derived from U^13^C glucose (1 g/L) tracing of *E. coli* MG1655 in M9 minimal media and *S. aureus* USA300 in RPMI1640. LC-MS analysis of ^13^C labeled heat-killed *E. coli*-derived cis-vaccenic acid and C17:0 cyclopropane fatty acid, and heat-killed *S. aureus*-derived *anteiso*-C15:0 branched chain fatty acid (**B**), and bacterial fatty acid-loaded acylcarnitines (**C**) from naïve or trained BMDMs rechallenged with ^13^C HKEC or HKSA for 24-hours in the presence of antimycin A (AA), or vehicle control (DMSO). **D.** LC-MS analysis of bacterial fatty acid recycling into the host lipidome compared between naïve or trained BMDMs stimulated with ^13^C HKEC or HKSA for 24 h in the presence of AA or vehicle control and measured as the Log_2_ fold change over naïve BMDMs treated with vehicle control and ^13^C HKEC of relative abundance of major isotopologues described in panel **A** of MS^2^ verified lipids within each lipid class: diacylglycerol (DG), triacylglycerol (TG), phosphatidylcholine (PC), phosphatidylethanolamine (PE), phosphatidylserine (PS), and sphingomyelin (SM). Graphs represent the mean of n ≥ 4 biological replicates. *p*-values were calculated with a two-way ANOVA with Sidak’s post-test. For **D**, statistics refer to trained + DMSO vs. trained + AA normalized peak area sums across validated species within each class. ns > 0.05; ∗*p* < 0.05; ∗∗*p* < 0.01; ∗∗∗*p* < 0.001; ∗∗∗∗*p* < 0.0001.

**Fig S8.**
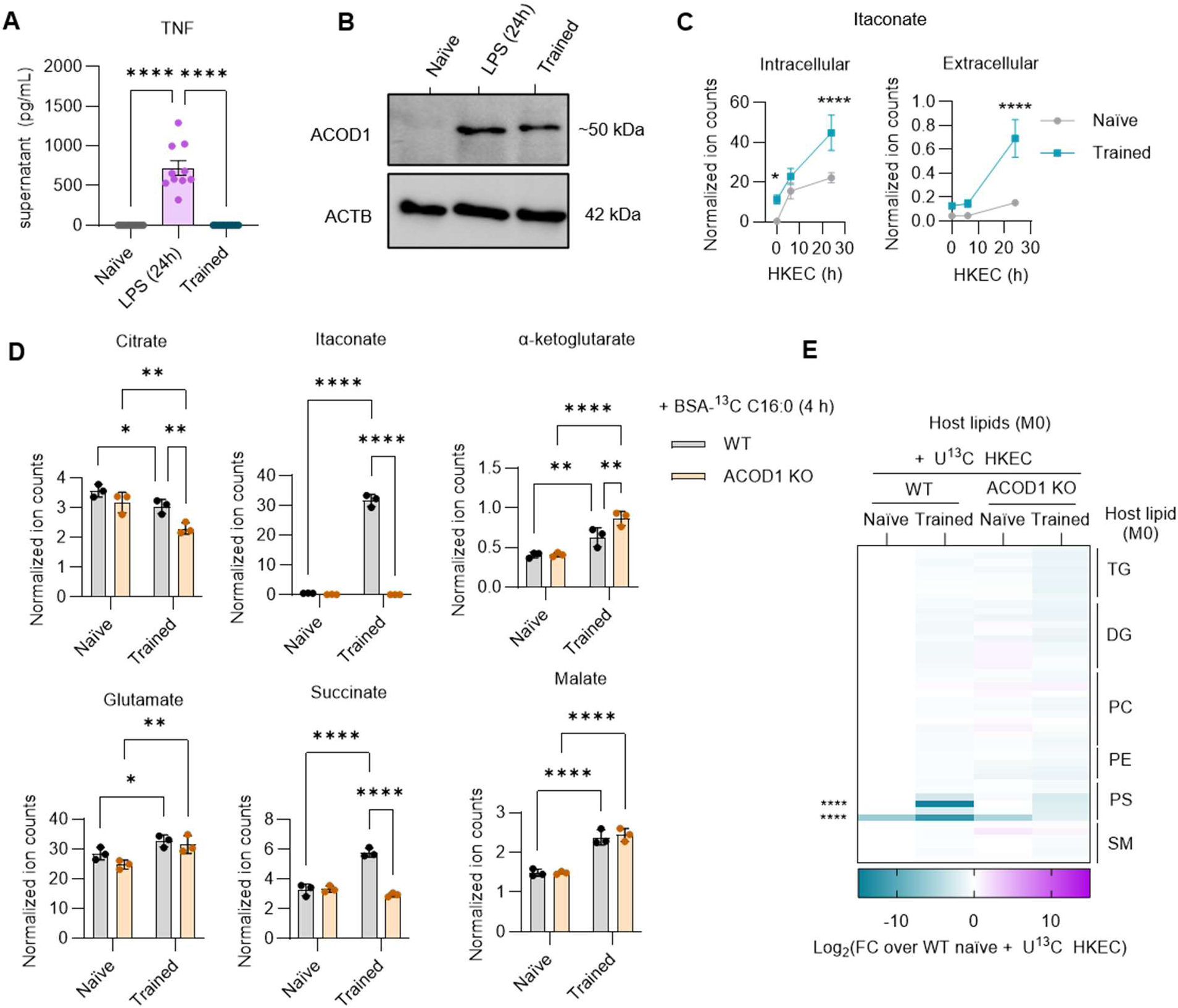
Host-derived metabolite and lipid abundances in trained ACOD1 KO macrophages. **A.** ELISA analysis of secreted TNF from naïve, continuous LPS stimulation (24 h), and LPS-trained (24 h after LPS wash-out) BMDMs’ supernatant. **B.** Western blot analysis of ACOD1 expression from lysates of naïve, LPS-stimulated, and LPS-trained BMDMs, generated as described in panel **A** with β-actin (ACTB) as a loading control. **C.** Detection of intracellular and extracellular itaconate produced by naïve and trained BMDMs stimulated with or without HKEC for 6 or 24 h. **D.** Pool size relative abundances of TCA cycle intermediates in naïve or trained WT and ACOD1 KO BMDMs, paired to analysis presented in **Fig 3D**. **E.** LC-MS analysis of relative abundances of indicated host lipids (M0), corresponding to lipid species highlighted in **Fig 3E** compared between naïve or trained BMDMs stimulated with ^13^C HKEC for 24 h in the presence of ETO, AA, or DMSO and measured as the Log_2_ fold change over naïve WT BMDMs treated with vehicle control and ^13^C HKEC of relative abundance of individual M+0 isotopologues of MS^2^ verified lipids within each lipid class, diacylglycerol (DG), triacylglycerol (TG), phosphatidylcholine (PC), phosphatidylethanolamine (PE), phosphatidylserine (PS), and sphingomyelin (SM). Graphs represent the mean of n ≥ 3 biological replicates with SEM error bars for all graphs. *p*-values were calculated with a one-way ANOVA with Tukey’s post-test (**A**) and two-way ANOVA with Sidak’s post-test (**C-E**). ns > 0.05; ∗*p* < 0.05; ∗∗*p* < 0.01; ∗∗∗*p* < 0.001; ∗∗∗∗*p* < 0.0001.

**Fig S9.**
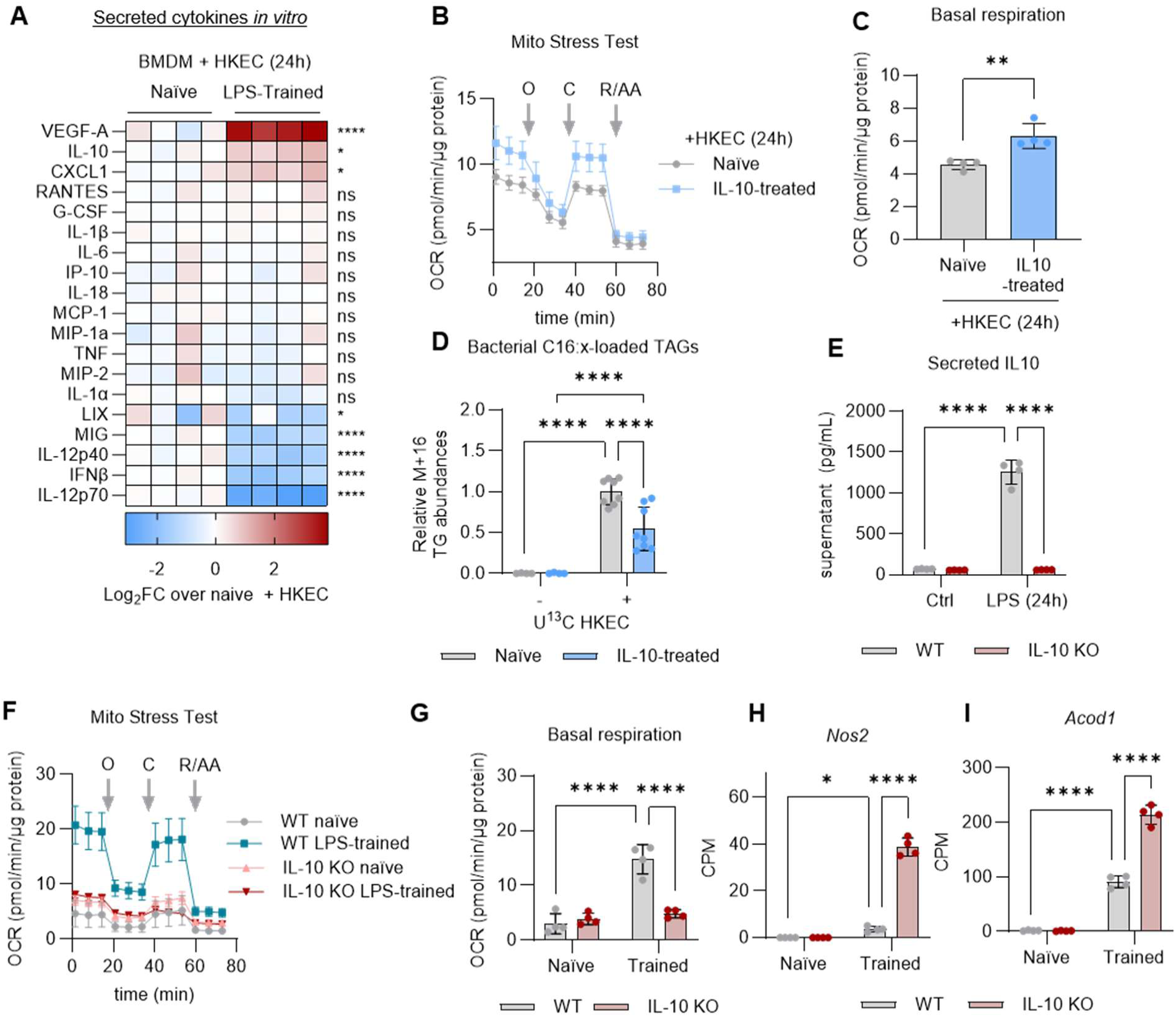
Investigation of IL-10 in sustaining OXPHOS in trained macrophages. **A.** Luminex analysis of secreted cytokines from naïve and trained BMDMs challenged with HKEC for 24h. Data are presented as the Log_2_ fold change over the average naïve + HKEC condition. **B.** Seahorse XF analysis of protein-normalized oxygen consumption rate (OCR) in naïve or IL-10 24h pretreated (IL-10-treated) BMDMs using the Mito Stress Test (MST) assay, with additions of oligomycin (O), Carbonyl cyanide m-chlorophenyl hydrazone (CCCP), rotenone (R), and antimycin A (A) at indicated time points. **C.** Quantification of basal respiration from panel **B. D.** LC-MS analysis of bacterial fatty acid recycling into the host TG pools (relative abundance of M+16 isotopologues) compared between naïve or IL-10-treated BMDMs fed ^13^C HKEC for 24h. **E.** ELISA analysis of secreted IL-10 levels from control or 24h LPS-stimulated WT and IL-10 KO BMDMs’ supernatant. **F.** Seahorse XF analysis of protein-normalized OCR in WT or IL-10 KO BMDMs using MST assay, as described above. **G.** Quantification of basal respiration from the MST assay. Highlighted RNASeq transcript-level analysis (counts per million; CPM) of *Nos2* (**H**) and *Acod1* (**I**). Graphs represent the mean of n = 4 biological replicates with SEM error bars for all graphs. *p*-values were calculated with a two-way ANOVA with Sidak’s post-test. ns > 0.05; ∗*p* < 0.05; ∗∗*p* < 0.01; ∗∗∗*p* < 0.001; ∗∗∗∗*p* < 0.0001.

## Notes

### Competing Interest Statement

The authors have declared no competing interest.

